# Evolution in action: Habitat-transition leads to genome-streamlining in Methylophilaceae (Betaproteobacteriales)

**DOI:** 10.1101/651331

**Authors:** Michaela M. Salcher, Daniel Schaefle, Melissa Kaspar, Stefan M. Neuenschwander, Rohit Ghai

## Abstract

The most abundant aquatic microbes are small in cell and genome size. Genome-streamlining theory predicts gene loss caused by evolutionary selection driven by environmental factors, favouring superior competitors for limiting resources. However, evolutionary histories of such abundant, genome-streamlined microbes remain largely unknown. Here we reconstruct the series of steps in the evolution of some of the most abundant genome-streamlined microbes in freshwaters (‘*Ca*. Methylopumilus’) and oceans (marine lineage OM43). A broad genomic spectrum is visible in the family Methylophilaceae (Betaproteobacteriales), from sediment microbes with medium-sized genomes (2-3 Mbp genome size), an occasionally blooming pelagic intermediate (1.7 Mbp), and the most reduced pelagic forms (1.3 Mbp). We show that a habitat transition from freshwater sediment to the relatively oligotrophic pelagial was accompanied by progressive gene loss and adaptive gains. Gene loss has mainly affected functions not necessarily required or advantageous in the pelagial or are encoded by redundant pathways. Likewise, we identified genes providing adaptations to oligotrophic conditions that have been transmitted horizontally from pelagic freshwater microbes. Remarkably, the secondary transition from the pelagial of lakes to the oceans required only slight modifications, i.e., adaptations to higher salinity, gained via horizontal gene transfer from indigenous microbes. Our study provides first genomic evidence of genome-reduction taking place during habitat transitions. In this regard, the family Methylophilaceae is an exceptional model for tracing the evolutionary history of genome-streamlining as such a collection of evolutionarily related microbes from different habitats is practically unknown for other similarly abundant microbes (e.g., ‘*Ca*. Pelagibacterales’, ‘*Ca*. Nanopelagicales’).

## Introduction

Marine and freshwater pelagic habitats are numerically dominated by very small microbes (cell volumes <0.1 μm^3^) that seem to be perfectly adapted to nutrient-poor (oligotrophic) conditions by successfully competing for dissolved organic matter and nutrients at low nM concentrations due to higher surface-to-volume ratios and superior transport systems [1]. Small-sized cells also enjoy other benefits such as reduced replication costs and mortality rates by size selective protistan predators [2]. The genomes of such oligotrophs are characterized by being very small (streamlined, <1.5 Mbp) with highly conserved core genomes and few pseudogenes, compacted intergenic spacers, reduced numbers of paralogs, and a low genomic GC content [3, 4]. While genetic drift has been proposed as the evolutionary mechanism behind the reduced genomes of symbionts, parasites and commensals, selection driven by environmental factors has been suggested as the primary driving force in the case of free-living oligotrophs [3]. The most abundant organisms on earth, bacteria of the marine SAR11 lineage (‘*Ca*. Pelagibacter ubique’, Alphaproteobacteria) serve as models for genome streamlining in the oceans [5] and their freshwater sister lineage LD12 is also known to be of similarly small size [6, 7]. Other examples of aquatic microbes with small cells and reduced genomes can be found among Actinobacteria (marine ‘*Ca*. Actinomarina minuta’ [8], freshwater ‘*Ca*. Nanopelagicales’ [9, 10], freshwater luna1 lineage [11, 12]), Thaumarchaeota (marine ‘*Ca.* Nitrosopelagicus brevis’)[13], and Betaproteobacteriales (freshwater ‘*Ca*. Methylopumilus planktonicus’ [14], marine OM43 lineage [15, 16]).

The latter are methylotrophs that are specialized in using reduced one-carbon (C_1_) compounds like methanol, methylamine and formaldehyde as sole energy and carbon sources by means of a modular system of different pathways for their oxidation, demethylation and assimilation [17]. The family Methylophilaceae (Betaproteobacteriales) is among the most important methylotrophs playing a key role in the carbon cycle of aquatic habitats [17, 18]. Four genera are so far validly described (*Methylotenera*, *Methylobacillus, Methylophilus, Methylovorus*) that mainly inhabit the sediment of freshwater lakes [19–22]. Axenic strains have been also isolated from the pelagial of lakes (‘*Ca*. Methylopumilus’)[14] and oceans (lineage OM43) [15, 16, 23]. Freshwater ‘*Ca*. Methylopumilus planktonicus’ are ubiquitous and highly abundant in lakes [24] with distinct maxima during diatom and/or cyanobacterial blooms [14, 25, 26], indicating that C_1_ substrates supporting their growth are released from primary producers. Members of the coastal marine OM43 lineage display similar temporal patterns with highest numbers during phytoplankton blooms [27–29].

In this work, we analysed the evolutionary history of the family Methylophilaceae by comparative genomics. While sediment dwellers have a larger cell and genome size, pelagic lineages are genome-streamlined. We hypothesize that the evolutionary origin of the family can be traced back to freshwater sediments, from where these microbes emerged to colonize the plankton of lakes and eventually also crossing the freshwater-marine boundary. The transition from sediments to the pelagial resulted in a pronounced genome reduction and adaptive gene loss has mainly affected functions that are not necessarily required or advantageous in the pelagial or are encoded in redundant pathways. Likewise, genes providing adaptations to oligotrophic conditions might have been transmitted horizontally from indigenous pelagic microbes.

## Material and Methods

### Isolation of planktonic freshwater methylotrophs

Novel strains of ‘*Ca*. Methylopumilus’ and other Methylophilaceae were isolated from the pelagial of Lake Zurich (CH), Římov Reservoir (CZ), and Lake Medard (CZ). Dilution-to-extinction using filtered (0.2 μm) and autoclaved water amended with vitamins and amino acids as a medium was used for Lake Zurich[14]. A full-cycle isolation approach [30] was employed for samples from Římov Reservoir and Lake Medard, with filtered water samples (0.45 μm filters), being diluted 1:10 with Artificial Lake Water (ALW [31]) containing vitamins (0.593 μM thiamine, 0.08 μM niacin, 0.000074 μM cobalamine, 0.005 μM para-amino benzoic acid, 0.074 μM pyridoxine, 0.081 μM pantothenic acid, 0.004 μM biotin, 0.004 μM folic acid, 0.555 μM myo-inosito, 0.01 μM riboflavin), 30 μM LaCl_3_, 1 mM methanol and 0.1 mM methylamine and incubated for 1-2 days at *in situ* temperatures. This step resulted in a preadaptation of methylotrophs only (C_1_ compounds as sole carbon source) without causing a shift in the assemblage of ‘*Ca*. Methylopumilus’ as these microbes displays slow growth with doubling times of approx. two days. Thereafter, a dilution to extinction technique was employed [14] with approx. 1 cell per cultivation well in 24-well-plates. Plates were incubated for 4-6 weeks at *in situ* temperature and growth in individual wells was checked microscopically and by PCR and Sanger sequencing of 16S rRNA genes.

### Whole-genome sequencing, assembly, and functional annotation

Thirty-eight pure cultures of ‘*Ca*. Methylopumilus sp.’ and three *Methylophilus* sp. were grown in 400 ml ALW medium supplemented with vitamins, LaCl_3_, methanol and methylamine for 6-8 weeks, pelleted by centrifugation, and DNA was isolated with a MagAttract® HMW DNA Kit (Qiagen). 550-bp libraries were constructed with the KAPA Hyper Prep Kit (Roche) and paired-end sequences (2 x 250-bp) were generated on an Illumina MiSeq instrument with a 500-cycle MiSeq Reagent v2 kit (Illumina). Library preparation and sequencing was done at the Genetic Diversity Center Zurich (GDC). Raw reads were quality trimmed with trimmomatic [32], assembled with SPAdes [33] and subsequently mapped to the resulting assemblies with Geneious 9 (www.geneious.com) in order to identify potential assembly errors]. Assembly usually resulted in 1-2 large contigs with overlapping ends that mostly could be circularized *in silico*. In the case of non-overlapping contigs, genomes were closed by designing specific primers for PCR and Sanger sequencing. Moreover, regions containing low coverage (≤10 fold), ambiguities, or anomalies in the mapping were verified by designing specific primers for PCR and Sanger sequencing to produce high-quality reference genomes. Gene prediction was done with PROKKA [34] and annotation was done with an in-house pipeline [10] based on BLAST searches to NCBI NR, COG [35], TIGRFAM [36] and KEGG databases [37]. Metabolic pathways were inferred from KEGG [37] and MetaCyc [38] and manually examined for completeness. Pathways involved in methylotrophy were identified by collecting 1016 reference protein sequences from published genomes of methylotrophs [14, 17, 39–45] and for the sake of completeness, also pathways not common to Methylophilaceae were included (e.g., methane oxidation [46–48], methylovory [49, 50]). These proteins were classified into 25 modules representing distinct (or sometimes alternative) biochemical transformations relevant to a methylotrophic lifestyle (e.g. M01-methanol oxidation, M02-pyrroloquinoline quinone biosynthesis, etc.; a complete list is provided in Supplementary Table S5). Protein sequences were clustered at 90% identity and 90% coverage with cd-hit [51] and the clusters were aligned using muscle [52]. The alignments were converted to HMMs (Hidden Markov Models) using the hmmbuild program in the HMMER3 package [53]. The program hmmsearch was used to scan complete genomes using these HMMs using e-value cut-off of 1e-3. The entire set of HMMs is available as Supplementary Data Set.

### Fragment recruitment from metagenomes

Publicly available metagenomes gained from freshwater sediments (n=131), the pelagial of lakes (n=345), rivers (n=43), estuaries, brackish and coastal oceanic sites (n=53) as well as open oceans (n=201) were used for fragment recruitment (see Table S2 for sampling sites and SRR accessions). rRNA sequences in genomes were identified with barrnap (http://www.vicbioinformatics.com/software.barrnap.shtml) and masked to avoid biases, and metagenomic reads were queried against the genomes using BLASTN [54] (cut-offs: length ≥50 bp, identity ≥95%, e-value ≤1e-5). These hits were used to compute RPKG values (number of reads recruited per kb of genome per Gb of metagenome), which provide a normalized value that is comparable across different genomes and metagenomes.

### Comparative genomic analyses

All publicly available genomes of high quality (>95% completeness, <20 scaffolds) affiliated with the family Methylophilaceae (Table S1, n=37) were downloaded from NCBI and re-annotated for comparative analyses. Average nucleotide identities (ANI [55]) and average amino acid identities (AAI [56]) were calculated to discriminate different species and genera. Phylogenomic trees based on conserved concatenated protein sequences (351,312 amino acid sites from 878 proteins for all Methylophilaceae, Fig. 1; 337,501 amino acid sites from 983 proteins for all ‘*Ca*. Methylopumilus’ spp., Fig. S1) was generated with FastTree [57] (100 bootstraps) after alignment with kalign [58]. *Methyloversatilis* sp. RAC08 (NZ_CP016448) and ‘*Ca*. Methylosemipumilus turicensis’ MMS-10A-171 (NZ_LN794158) served as outgroup for the trees displayed in Fig. 1 and Fig. S1, respectively. The core- and pangenome of the family was computed using all-vs-all comparisons of all proteins for each genome using BLASTP (≥50% identity and ≥50% coverage cut-offs to define an ortholog). Paralogs in each genome were identified with BLASTP (cut-offs: ≥80% coverage, ≥70% similarity, ≥50% identity). Closest relatives for proteins putatively transferred horizontally were identified with BLASTP against the NCBI Protein Reference Sequences database (cut-off: *E* values ≤1e-5). Trees for individual proteins or concatenated proteins for specific pathways were constructed with RAxML (GAMMA BLOSUM62 model [59]) after alignment with MAFFT v7.388 [60].

**Figure 1:**
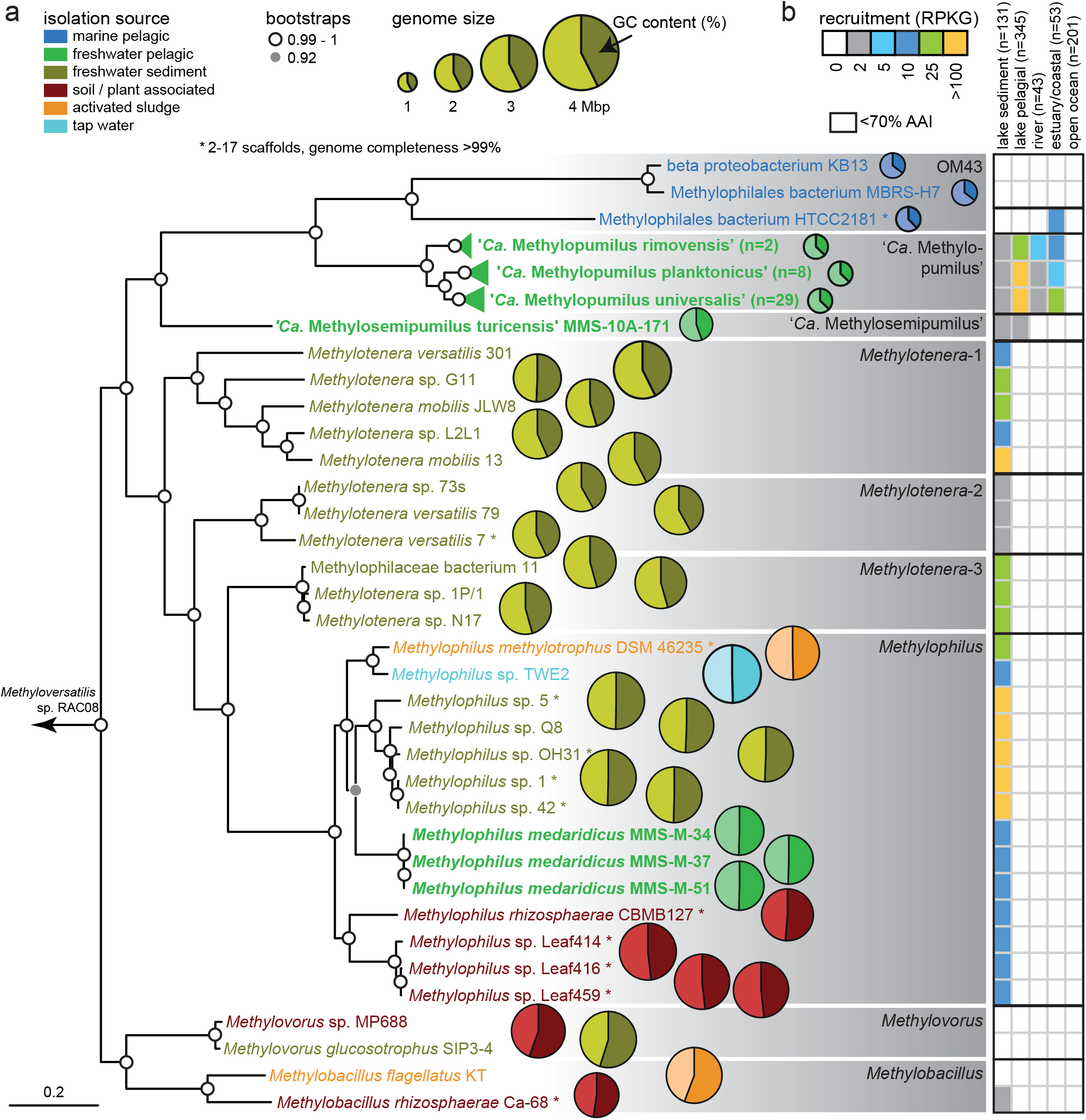
Phylogeny of Methylophilaceae and their occurrence in different environments. (a) Phylogenomic tree based on 878 common concatenated proteins (351,312 amino acid sites) with *Methyloversatilis* sp. RAC08 as outgroup. The 39 complete genomes of ‘*Ca*. Methylopumilus sp.’ are collapsed to the species level, see Fig. S1 for a complete tree. Different genera (70% AAI cut-off, Fig. S2) are marked by grey boxes. Isolation sources of strains are indicated by different colours and incomplete genomes consisting of <17 contigs (estimated completeness >99%) are marked with asterisks. Bootstrap values (100 repetitions) are indicated at the nodes, the scale bar at the bottom indicates 20% sequence divergence. The genome sizes for all strains are shown with circles of proportional size and GC content is depicted within each circle. (b) Fragment recruitment of public metagenomes from freshwater sediments (n=131), lake pelagial (n=345), rivers (n=43), estuaries and coastal oceans (n=53), and open oceans (n=201). Maximum RPKG values (number of reads recruited per kb of genome per Gb of metagenome) for each ecosystem are shown for each genome.

### Availability of data

All genomes have been submitted to NCBI under BioProject XXX, BioSamples XY-XYX (Please note: Submission is in progress, accession numbers will be provided as soon as possible). The entire set of HMMs related to methylotrophic functions is available as Supplementary Data Set.

## Results and discussion

### Phylogenomics and global occurrence of Methylophilaceae

Currently, 31 Methylophilaceae genomes of high quality (i.e., >99% completeness, <20 scaffolds) are publicly available, mostly from axenic isolates from freshwater sediments (Fig. 1, Table S1). We additionally sequenced the genomes of 41 strains of planktonic freshwater strains affiliated with ‘*Ca*. Methylopumilus planktonicus’ (38 strains) and *Methylophilus* sp. (3 strains). These microbes were isolated from the pelagial of three different freshwater habitats (Lake Zurich, CH; Římov Reservoir, CZ; Lake Medard, CZ) by dilution-to-extinction [14, 30]. All novel genomes are of very high quality, i.e., they are complete, with one circular chromosome (Table S1). The 39 strains classified as ‘*Ca*. M. planktonicus’ by 16S rRNA gene sequences analysis (99.94-100% sequence identity), constitute at least three different species according to average nucleotide and amino acid identity (ANI and AAI [61]) (Fig. S1-S4). We tentatively name these three taxa ‘*Ca*. Methylopumilus rimovensis’ (two strains isolated from Římov Reservoir), ‘*Ca*. Methylopumilus universalis’ (29 strains from Lake Zurich and Římov Reservoir) and the originally described ‘*Ca*. Methylopumilus planktonicus’ (eight strains from Lake Zurich; Fig. 1, Fig. S1)[14]. AAI values also suggest that ‘*Ca*. M. turicensis’ is a different genus (62% AAI with ‘*Ca*. Methylopumilus’, Fig. S3) that we tentatively rename to ‘*Ca*. Methylosemipumilus turicensis’. This reclassification is in line with the recently released Genome Taxonomy Database (GTDB [62]). Moreover, the genus *Methylotenera* might be split in different genera and the GTDB suggests a reclassification of several strains to the genus *Methylophilus*. Our analysis notes a polyphyletic pattern of *Methylotenera* with three different genera (*Methylotenera*-1, *Methylotenera*-2, and *Methylotenera*-3; Fig. 1a, Fig. S3, >70% AAI). However, further work is necessary to clarify the formal naming of these strains as AAI values are inconclusive and the proposed hard cut-off of 65% AAI for genus delineation are not met for most members of Methylophilaceae. Three novel pelagic *Methylophilus* sp. isolates (MMS-M-34, MMS-M-37, MMS-M-51) constitute a novel species that we tentatively named *M. medardicus*, with closest hits to isolates from freshwater sediment. These strains might originate from the same clone, as they were gained from the same sample from Lake Medard and were 100% identical in in their genome sequence. *M. mediardicus* seem to be not abundant in the pelagial of lakes, as indicated by recruitments from 345 different pelagic freshwater metagenomic datasets, however they could be readily detected in relatively high proportions in sediment metagenomes (Fig. 1b, Table S2). Sediments also appear to be the main habitat of other *Methylophilus* and *Methylotenera*. The three strains isolated from marine systems, that were referred to as OM43-lineage [15, 16, 23], form two different genera based on AAI (Fig. S3). However, none appear to be abundant in the open ocean (Fig. 1b), and only strain HTCC2181 could be detected in estuarine/coastal metagenomes, although lineage OM43 has been repeatedly reported in coastal oceans by CARD-FISH, where they can reach up to 4% or 0.8 x 10^5^ cells ml^−1^ during phytoplankton blooms [28, 29]. It is thus likely that other, more abundant strains of OM43 still await isolation. ‘*Ca*. Methylopumilus spp.’ on the other hand, were found in moderate proportions in estuarine/coastal systems, but their main habitat is clearly the pelagial of lakes, where they are highly abundant (Fig. 1b), as previously reported based on CARD-FISH analyses [14, 63], 16S rRNA gene amplicon sequencing [24, 64–66], and metagenomics [67, 68]. All ‘*Ca*. Methylopumilus’, especially ‘*Ca*. M. rimovensis’ were also prevalent in rivers (Fig. 1b).

### Genome-streamlining in pelagic strains

The genomes of pelagic freshwater ‘*Ca*. Methylopumilus sp.’ (n=39) and the marine OM43 lineage (n=3) are characterized by very small sizes (1.26-1.36 Mbp) and a low genomic GC content (35.3-37.7%) (Table S1, Fig. 2, Fig. S5). ‘*Ca*. Methylosemipumilus turicensis’ MMS-10A-171 has a slightly larger genome (1.75 Mbp) with higher GC content (44.5%), while all other Methylophilaceae have genome sizes >2.37 Mbp (max. 3.25 Mbp) and a higher GC content (41.9-55.7%, average 47.3%). A highly significant relationship between genome size and GC content, length of intergenic spacers, coding density, mean CDS length, number of overlapping CDS, paralogs, and numbers of genes involved in sensing of the environment (i.e., histidine kinases and sigma factors) was evident (Fig. 2). All these features have been proposed to be relevant for genome-streamlining [3] with freshwater ‘*Ca*. Methylopumilus’ and marine OM43 displaying the most reduced forms and ‘*Ca*. Methylosemipumilus turicensis’ presenting an intermediate state (Table S1). Moreover, we observed a negative relationship between genomic GC content and stop-codon usage of TAA instead of TAG, as well as a preferred amino acid usage of lysine instead of arginine (Fig. 2), both suggested to be involved in nitrogen limitation [4]. Furthermore, amino acids with less nitrogen and sulphur and more carbon atoms were favourably encoded by the genome-streamlined microbes (Fig. 2, S5, S6).

**Figure 2:**
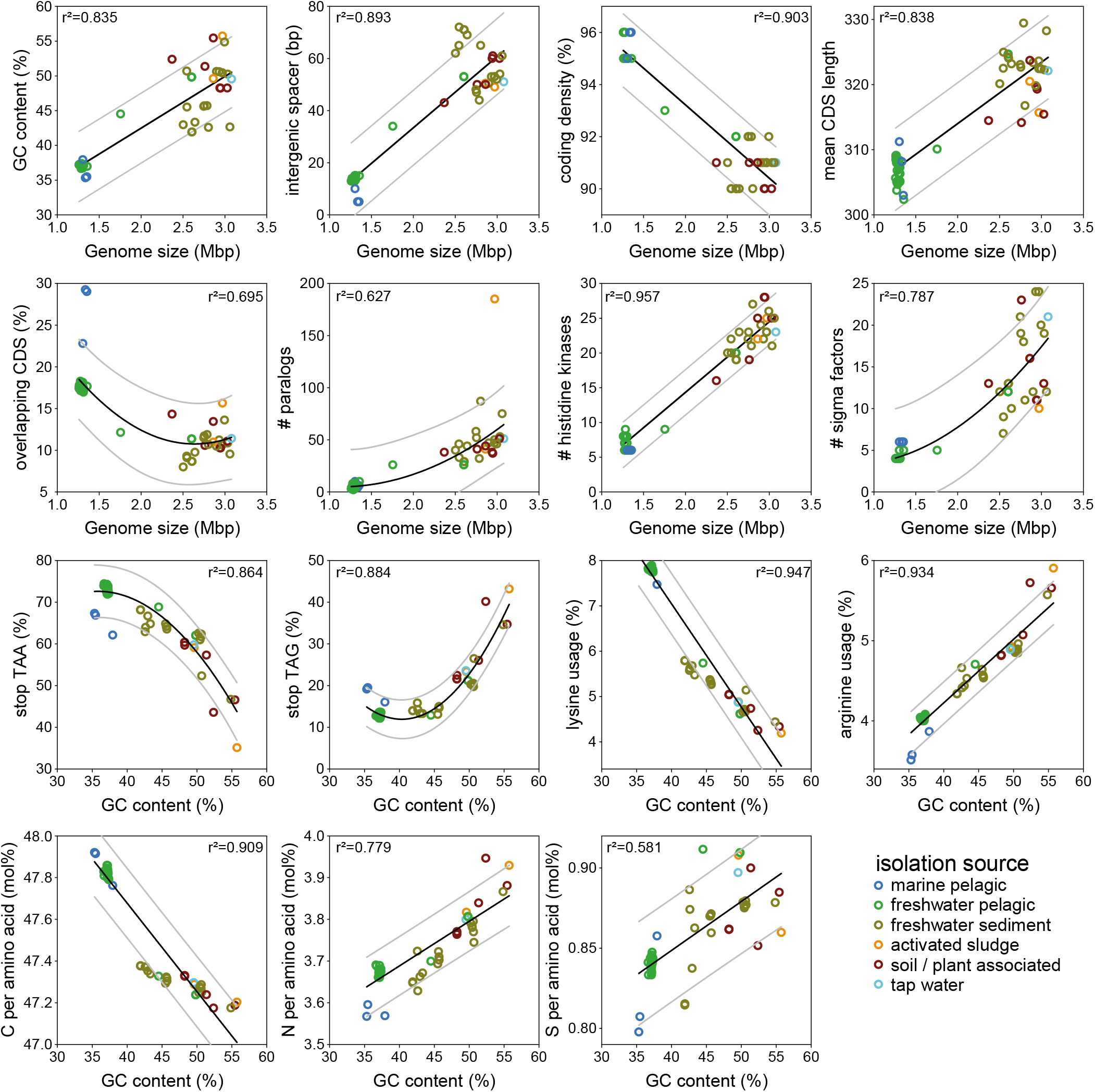
Genome streamlining in Methylophilaceae. Significant relationships between genome sizes (Mbp) and genomic GC content (%), lengths of intergenic spacers (bp), coding density (%), mean CDS length (bp), overlapping CDS (%), number of paralogs, histidine kinases and sigma factors and significant relationships between genomic GC content (%) and TAA stop codon, TAG stop codon, lysine, and arginine usage (%) and C, N, and S atoms per amino acid (mol%).

### Adaptive gene loss during habitat transition from the sediment to the pelagial

The core genome of the family Methylophilaceae consists of 664 protein families (4.3% of the pangenome) and an open pangenome of >15,000 protein families, while the streamlined genomes of ‘*Ca*. Methylopumilus’ have a highly conserved core (48%, Fig. S7). By contrast, sediment Methylophilaceae have a larger pangenome with a more modular assortment featuring several redundant pathways for methylotrophic functions [18] and a large fraction of proteins overlapping with ‘*Ca*. M. turicensis’, indicating a high evolutionary relatedness (Fig. S7). It appears that both extant pelagic and sediment Methylophilaceae shared a common sediment-dwelling methylotrophic ancestor. While one lineage (*Methylotenera* and *Methylophilus*) retains the ancestral character (large genomes) of the common ancestor, the other lineage diversified towards a pelagic lifestyle (‘*Ca*. M. turicensis’, ‘*Ca*. Methylopumilus’ and OM43). Remarkably, ‘*Ca*. M. turicensis’ appears to constitute an early diverging lineage that displays somewhat mixed characteristics of both sediment and truly pelagic forms (‘*Ca*. Methylopumilus’ and OM43), not only in its phylogenetic position, but also in genomic characteristics. Monitoring data from Lake Zurich showed consistently high cell densities of ‘*Ca*. Methylopumilus’, while ‘*Ca*. M. turicensis’ were mostly below detection limits except for a 3-month phase in one year where they reached high numbers in the hypolimnion [14]. Moreover, also fragment recruitment from freshwater metagenomes showed a global occurrence of ‘*Ca*. Methylopumilus’ in very high relative proportions, while ‘*Ca*. M. turicensis’ was less prevalent (Fig. 1b). This hints again at the somewhat transitional character of ‘*Ca*. M. turicensis’ (occasional ‘bloomer’) that is not as perfectly adapted to the pelagial as ‘*Ca*. Methylopumilus’.

We tested the hypothesis of adaptive gene loss driven by evolutionary selection during the transition from sediment to the pelagial by comparative genomics of metabolic modules of *Methylophilus*, *Methylotenera*, ‘*Ca*. M. turicensis’, ‘*Ca*. Methylopumilus’ and marine OM43 strains. *Methylobacillus* and *Methylovorus* were excluded as they seem too distantly related and also not very abundant in lake sediments (Fig. 1b, Table S2). The most pronounced differences in the genetic make-up of sediment vs. pelagic strains were detected in motility and chemotaxis (Figs. 3, 4), with all *Methylophilus* and all but two *Methylotenera* strains having flagella and type IV pili, while the planktonic strains have lost mobility and also greatly reduced the number of two-component regulatory systems and sigma factors. A large number of membrane transporters for inorganic compounds was detected exclusively in sediment Methylophilaceae, while this number is reduced in ‘*Ca*. M. turicensis’ and even more in ‘*Ca*. Methylopumilus’ and OM43 (Fig. 4, Table S3). Moreover, *Methylophilus* and *Methylotenera* encode multiple pathways for nitrogen acquisition, with transporters for ammonia, nitrate, nitrite, taurine, cyanate or urea, and pathways for urea or cyanate utilization (Fig. 3, Table S4)[69]. ‘*Ca*. M. turicensis’, on the other hand, has only transporters for nitrate/taurine and ammonia (Amt family), and ‘*Ca*. Methylopumilus’ and marine OM43 only carry ammonia transporters. All sediment Methylophilaceae further possess genes for assimilatory nitrate reduction to ammonia, some for dissimilatory nitrate reduction to nitrous oxide or detoxification of nitric oxide (quinol type of *norB*) [70] and *Methylotenera mobilis* 13 is a complete denitrifier [69, 71, 72], while none of the pelagic strains have any genes involved in nitrate reduction (Figs. 3, 4, Table S4). Ammonia is the main microbial nitrogen source in the epilimnion of lakes and oceans, while nitrate and other compounds like urea, taurine, or cyanate are more abundant in deeper, oxygenated layers and the sediment [14, 73, 74]. Therefore, an adaptation to ammonium uptake might be advantageous for pelagic microbes.

**Fig. 3:**
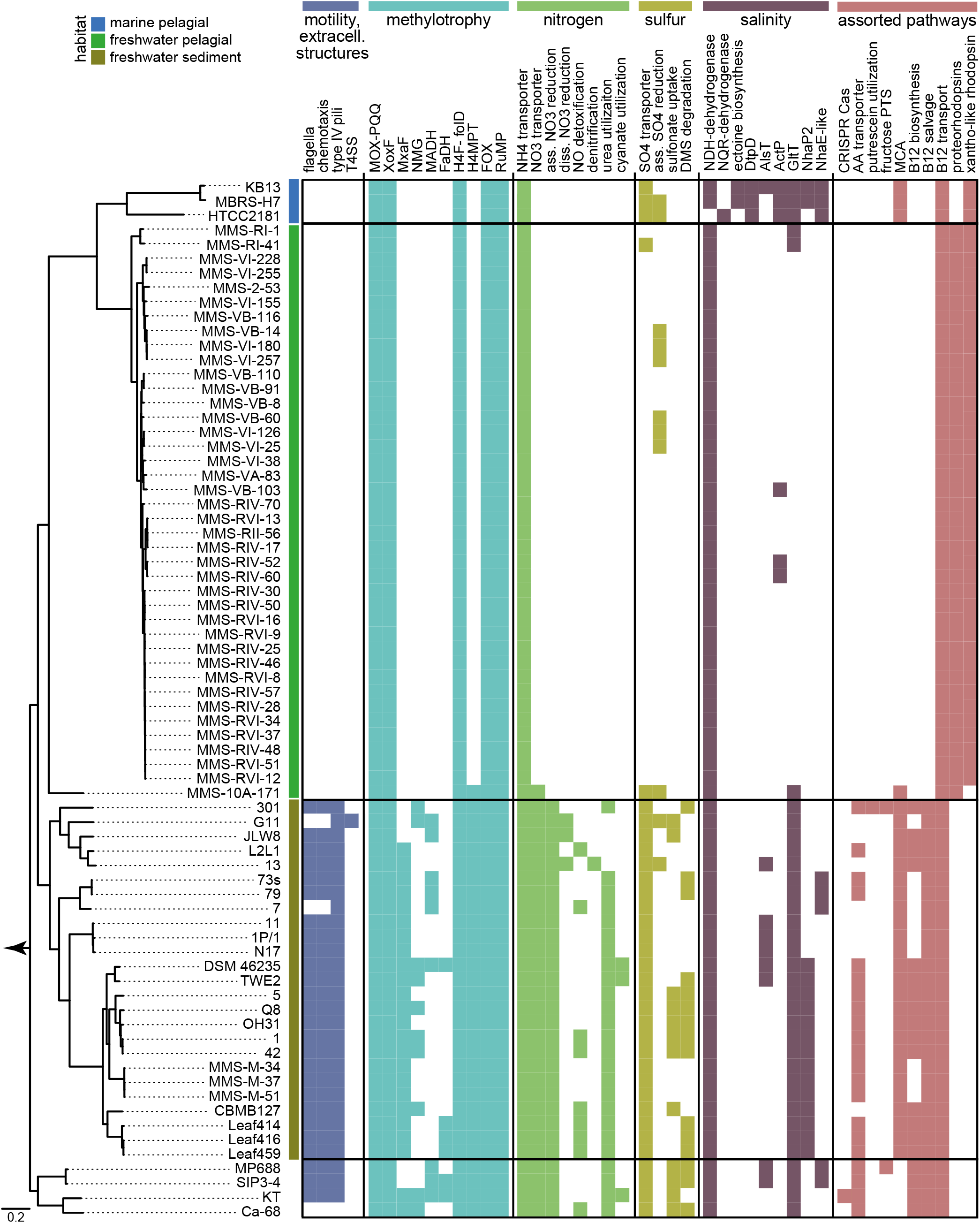
Metabolic modules in Methylophilaceae. Presence of selected metabolic modules in Methylophilaceae strains. For details on phylogenomic tree see Fig. 1 and Fig. S1, for details on pathways see Table S4. T4SS: type 4 secretion system; MOX-PQQ: methanol oxidation and pyrroloquinoline quinone biosynthesis; XoxF/MxaF: methanol dehydrogenase XoxF/MxaF; NMG: N-methyloglutamate pathway; MADH: methylamine dehydrogenase; FaDH: formaldehyde dehydrogenase; H4F-folD: H_4_-linked formaldehyde oxidation, folD form; H4MPT: H_4_MPT-linked formaldehyde oxidation; FOX: formate oxidation; RuMP: ribulose monophosphate cycle; ass. NO3 reduction: assimilatory nitrate reduction; diss. NO3 reduction: dissimilatory nitrate reduction; ass. SO4 reduction: assimilatory sulfate reduction; NDH-dehydrogenase: H^+^-translocating NADH:quinone oxidoreductase; NQR-dehydrogenase: Na^+^-translocating NADH:quinone oxidoreductase; DtpD: dipeptide/tripeptide permease DtpD; AlsT: sodium:alanine symporter AlsT; ActP: sodium:acetate symporter ActP; GltT: sodium:dicarboxylate symporter GltT; NhaP2: sodium:proton antiporter NhaP2; NhaE-like: sodium:proton antiporter NhaE; AA transporter: amino acid transporters; fructose PTS: fructose-specific phosophotransferase system; MCA: methylcitric acid cycle.

**Figure 4:**
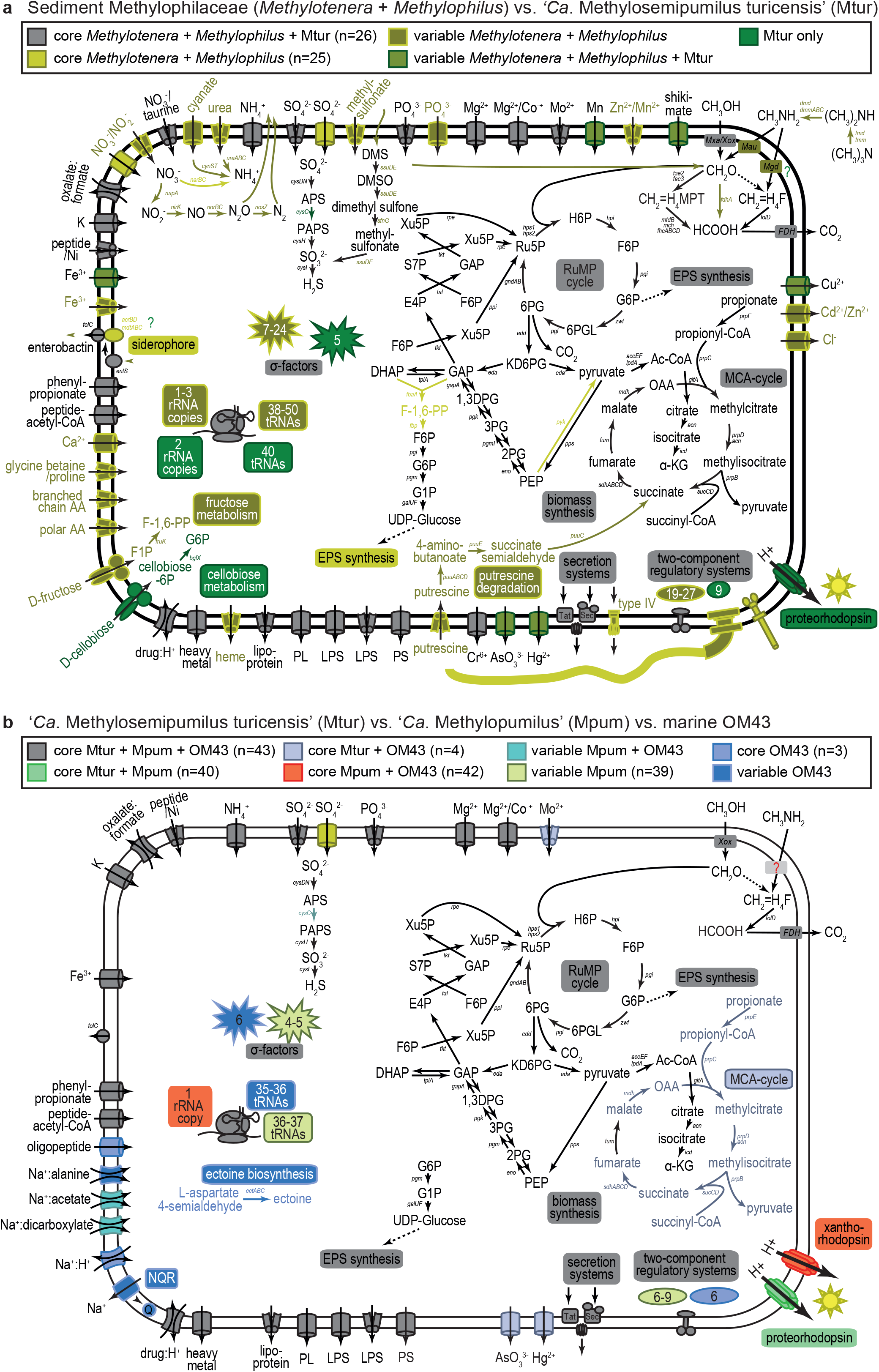
Comparative metabolic maps of different taxa of Methylophilaceae. (a) Comparison of the core metabolism in sediment Methylophilaceae (*Methylotenera* and *Methylophilus*) vs. ‘*Ca*. Methylosemipumilus turicensis’ (Mtur). (b) Comparison of the core metabolism in ‘*Ca*. Methylosemipumilus turicensis’ (Mtur) vs. ‘*Ca*. Methylopumilus’ (Mpum) vs. marine OM43. For details on pathways see Table S4.

Furthermore, a high diversity of pathways involved in sulfur metabolism was detected in Methylophilaceae, with the genome-streamlined strains representing the most reduced forms again. All *Methylophilus*, *Methylotenera,* ‘*Ca*. Methylosemipumilus turicensis’ and one strain of ‘*Ca*. Methylopumilus rimovensis’ encode ABC transporters for sulfate uptake, and a sulfate permease was annotated in OM43 and several sediment Methylophilaceae, while the majority of ‘*Ca*. Methylopumilus’ lack these transporters (Fig. 3, 4). Canonical assimilatory sulfate reduction seems to be incomplete in most Methylophilaceae, as adenylyl sulfate kinase *cysC* was annotated only in a few strains (Table S4). Thus, the mode of sulfite generation remains unclear, with unknown APS kinases or other links from APS to sulfite. *Methylophilus rhizosphaera* encodes genes for dissimilatory sulfate reduction and most sediment strains possess ABC transporters for alkanesulfonates, most likely transporting methylsulfonate, that can be oxidised to sulfite by methanesulfonate monooxygenases generating formaldehyde as by-product. Dimethyl sulfide (DMS) seems to be a source for sulfur and formaldehyde as well, as dimethylsulfoxide and dimethyl monooxygenases are present in several sediment Methylophilaceae, but absent in all pelagic strains. It is thus still unclear how ‘*Ca*. Methylopumilus’ fuel their sulfur demand, especially as they grow in a defined medium containing sulfate (200 μM MgSO_4_; 160 μM CaSO_4_) and vitamins as sole sulfur sources.

All Methylophilaceae have complete pathways for the biosynthesis of amino acids and vitamins, with the exception of cobalamin (vitamin B12) that was lacking in the pelagic lineage (‘*Ca*. M. turicensis’, ‘*Ca*. Methylopumilus’, marine OM43), while either the complete biosynthesis or the salvage pathway was present in the sediment isolates (Figs. 3, 4, Table S4). However, putative cobalamin transporters were annotated in all isolates.

The methylcitric acid (MCA) cycle for oxidising propionate via methylcitrate to pyruvate is present in *Methylotenera*, *Methylophilus*, ‘*Ca*. M. turicensis’, and the marine OM43, but absent in all ‘*Ca*. Methylopumilus’ strains, suggesting it has been selectively lost in these organisms. All genes were arranged in a highly conserved fashion, with the exception of ‘*Ca*. M. turicensis’ having a bifunctional aconitate hydratase 2/2-methylisocitrate dehydratase (*prpD/acnB*) gene and *acnB* genes being located in different genomic regions in OM43 and ‘*Ca*. M. turicensis’ (possessing two copies), however, with high synteny of flanking genes (Fig. S8). Phylogenetic analysis of the MCA gene cluster resulted in genus-specific branching, and notably, the MCA pathway of OM43 is most closely related to that of ‘*Ca*. M. turicensis’ (Fig. S8), suggesting it was retained in the OM43 lineage after divergence from a common ancestor of OM43 and ‘*Ca*. M. turicensis’.

### Genome-streamlining leading to a loss of redundant methylotrophic pathways

Some of the sediment dwellers seem to be facultative methylotrophs, as ABC transporters for amino acids were annotated (Figs. 3, 4, Table S3). *Methylotenera versatilis* 301 additionally encodes a fructose-specific phosphotransferase system (PTS) and a 1-phosphofructokinase, as well as transporters for putrescine uptake and the subsequent pathway for its degradation. ‘*Ca*. M. turicensis’ might also be a facultative methylotroph, as it possesses a PTS system for cellobiose, while this (as well as amino acid transporters) is lacking in all ‘*Ca*. Methylopumilus’ and OM43 strains, making them obligate methylotrophs. These observations suggest that the ancestor of both pelagic and sediment lineages was also a facultative methylotroph and that obligate methylotrophy emerged only in the truly pelagic strains.

Remarkably, also pathways involved in methylotrophy were reduced in the course of genome streamlining with the sediment dwelling *Methylophilus* and *Methylotenera* having the most complete modules for C_1_ compound oxidation, demethylation and assimilation (Fig. 3, 4, Table S4). They also encode multiple types of methanol dehydrogenases (up to five different types in single strains), while the pelagic forms possess only XoxF4-1 (Fig. S9). Moreover, the latter encode neither traditional methylamine-dehydrogenases nor the N-methylglutamate (NMG) pathway for methylamine oxidation. Thus, the mode of methylamine uptake is still unclear, although it has been experimentally demonstrated that some pelagic strains can utilize this C_1_ substrate [14, 75]. However, also nearly half of the sediment strains lack these well-described pathways in a patchy manner only partly reflected by phylogeny, therefore it is likely that methylamine utilization is not a common feature within Methylophilaceae, or that alternative routes of its oxidation still await discovery [69]. Formaldehyde oxidation can be achieved via three alternative routes, and only four *Methylophilus* strains encode all of them, i.e., all others lack a formaldehyde-dehydrogenase. All *Methylophilus* and *Methylotenera* as well as ‘*Ca*. M. turicensis’ carry genes for the tetrahydromethanopterin (H_4_MPT) pathway, but none of the ‘*Ca*. Methylopumilus’ and OM43 strains. Therefore, the only route for formaldehyde oxidation in these genome-streamlined microbes is the tetrahydrofolate (H_4_F) pathway which includes the spontaneous reaction of formaldehyde to H_4_F and is thought to be relatively slow [14, 17]. The ribulose monophosphate (RuMP) cycle for formaldehyde assimilation/oxidation and formate oxidation via formate dehydrogenases was annotated in all Methylophilaceae, while none of them possess other potential methylotrophic modules such as the serine cycle, the ethylmalonyl-CoA-pathway for glyoxylate regeneration, a glyoxylate shunt, nor the Calvin-Benson-Bassham cycle for CO_2_ assimilation, as already previously noted to be lacking in Methylophilaceae [17]. Thus, the core methylotrophic modules in Methylophilaceae contain methanol oxidation via XoxF methanol dehydrogenases, formaldehyde oxidation via the H_4_F pathway, the RuMP cycle, and formate oxidation (Fig. 3, Table S4)[17, 69, 76]. The majority of genes encoding these pathways were organized in operon structures or found in close vicinity to each other with high synteny and phylogenetically reflecting the overall phylogeny of the family (Fig. S10, S11).

### Photoheterotrophy as adaptation to oligotrophic pelagic conditions

Rhodopsins are light-driven proton pumps producing ATP that fuel e.g., membrane transporters [77] and play important roles during carbon starvation [78] in oligotrophic aquatic environments. Therefore, the acquisition of rhodopsins are proposed to be powerful adaptations to the pelagial. ‘*Ca*. Methylosemipumilus turicensis’ acquired a proteorhodopsin and the complete pathway for retinal biosynthesis via horizontal gene transfer (HGT) from the abundant freshwater microbe *Polynucleobacter cosmopolitanus* (81% amino acid similarity, Fig. 5, Fig. S12a). Interestingly, ‘*Ca*. Methylopumilus spp.’ carry, in addition to a proteorhodopsin highly similar to ‘*Ca*. M. turicensis’ (78.3-79.5% similarity), a second rhodopsin gene inserted between the proteorhodopsin and the retinal biosynthesis cluster (Fig. 5). This xantho-like rhodopsin was most likely gained from rare freshwater Betaproteobacteria (*Janthinobacterium lividum, Massilia psychrophila;* 49.4-53.5% similarity, Fig. S12b), however, the binding of a carotenoid antenna seems unlikely due to the replacement of a glycine with tryptophan in position 156, suggesting it also functions as a proteorhodopsin (Fig. S12c)[79, 80]. Both rhodopsins are tuned to green light, which is common in freshwaters [81] and possess the canonical DxxxK retinal binding motif in helix-7 that is characteristic of proton pumping rhodopsins [82]. The marine OM43 lineage only carry the xantho-like rhodopsin (59.0-63.9% similarity to ‘*Ca*. Methylopumilus’). It is unclear if the proteorhodopsin was never present in the marine lineage or was lost subsequently, and if so, the reasons for a secondary loss remain enigmatic as two rhodopsins would provide an even better adaptation to oligotrophic waters than one.

**Figure 5:**
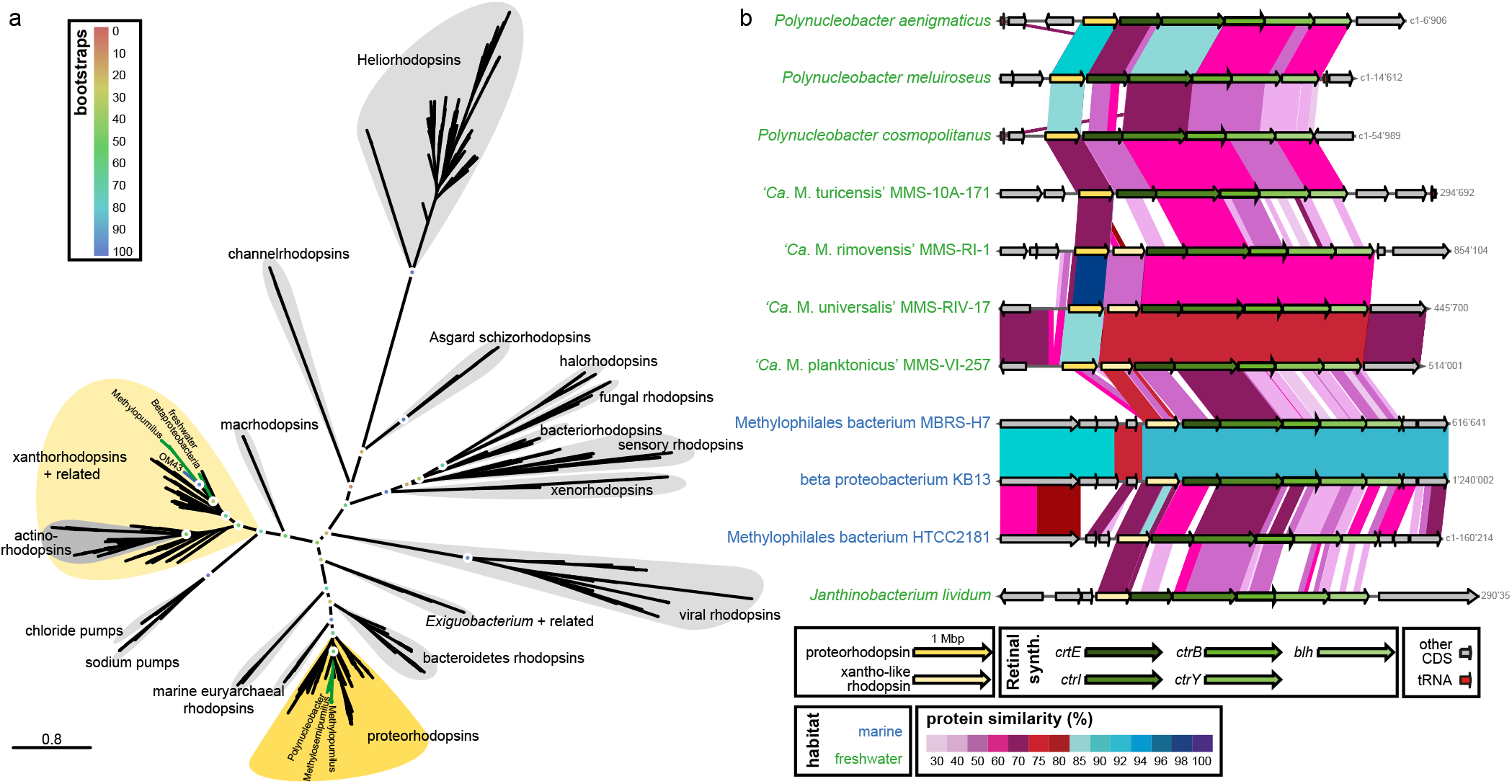
Horizontal gene transfers of two different rhodopsins. (a) Phylogenetic tree (RAxML, 100 bootstraps) of different rhodopsin types. See Fig. S12 for details of closely related proteo- and xantho-like rhodopsins of Methylophilaceae. (b) Arrangement and protein similarity of rhodopsins and the retinal biosynthesis gene cluster in ‘*Ca*. Methylopumilus spp.’, ‘*Ca*. Methylosemipumilus turicensis’, marine OM43 and other freshwater microbes with closely related rhodopsin types.

### The second transition from freshwater pelagial to the marine realm is characterized by adaptations to a salty environment

The second habitat transition across the freshwater-marine boundary does not appear to involve genome streamlining, as genomes of pelagic freshwater and marine methylotrophs are of similar small size and low GC content (Figs. 1, 2). We hypothesize that this transition had less impact on the lifestyle (purely planktonic, oligotrophic) but required specific adaptations to the marine realm that were mainly acquired by HGT, and as suggested by the long branches in the phylogenetic tree (Fig. 1), multiple, rapid changes in existing genes. Besides the MCA pathway and the proteorhodopsin, no major rearrangements or reductions in general metabolic pathways were detected in marine OM43 in comparison to freshwater ‘*Ca*. Methylopumilus’. However, several adaptations to higher salt concentrations could be identified (Figs. 3, 4). Salinity is one of the most important obstacle in freshwater-marine colonization, and successful transitions have occurred rarely during the evolution of Proteobacteria [83, 84]. Main adaptations to higher salinities involve genes for osmoregulation and inorganic ion metabolism that might have been acquired from the indigenous community by HGT. For example, genes regulating the Na^+^-dependent respiratory chain (Na^+^-translocating NADH:quinone oxidoreductase, NQR, Fig. S13) have been transmitted from the marine *Roseobacter* lineage to strain HTCC2181 [16, 84]. The NQR system provides energy by generating a sodium motif force, yet, the sodium pumping might also be an adaptation to enhanced salinities [85]. All other Methylophilaceae possess the energetically more efficient H^+^-translocating type (NDH), which works better under low salinity conditions and is thus common in freshwater microbes [85].

Ectoine, a compatible solute along with glycine betaine, helps organisms survive extreme osmotic stress by acting as an osmolyte [86]. Ectoine is synthesized from L-aspartate 4-semialdehyde, the central intermediate in the synthesis of amino acids of the aspartate family. Two marine OM43 strains (KB13 and MBRS-H7) encode this pathway followed by sodium:proline symporter *putP* arranged in high synteny and protein similarity with marine/hypersaline sediment microbes, thus it is likely that both components were gained via HGT (Fig. S14). A second copy of the *putP* symporter was common to all Methylophilaceae (data not shown). Also a dipeptide/tripeptide permease (DtpD) unique for the marine OM43 lineage seems to be transferred horizontally, either from marine Bacteroidetes or sediment-dwelling *Sulfurifustis* (Gammaproteobacteria, Fig. S15). Other putative membrane compounds involved in sodium transport in marine OM43 include a sodium:alanine symporter (AlsT, Fig. S16a), a sodium:acetate symporter (ActP, Fig. S16b), a sodium:dicarboxylate symporter (GltT, Fig. S16c), a sodium:proton antiporter (NhaP, Fig. S16d), and another putative sodium:proton antiporter (NhaE-like, Fig. S16e). Although also several other Methylophilaceae carry some of these sodium transporters, they are only distantly related to OM43, thus they might be acquired horizontally. Conversely, ActP and GltT of OM43 are most closely related to three ‘*Ca*. M. universalis’ strains and the two ‘*Ca*. M. rimovensis’ strains, respectively (Fig. S16b, S16c). Both symporters are related to microbes from freshwater and marine habitats, hinting to some yet unknown lineages related to both OM43 and ‘*Ca*. Methylopumilus’ most likely thriving in the freshwater-marine transition zone.

## Conclusions

Our study provides first genomic evidence that the ancestors of genome-streamlined pelagic Methylophilaceae can be traced back to sediments with two habitat transitions occurring in the evolutionary history of the family. The first from sediments to the pelagial is characterized by pronounced genome reduction driven by selection pressure for relatively more oligotrophic environmental conditions. This adaptive gene loss has mainly affected functions that (i) are not necessarily required in the pelagial (e.g., motility, chemotaxis), (ii) are not advantageous for survival in an oligotrophic habitat (e.g., low substrate affinity transporters), and (iii) are encoded in redundant pathways (e.g., formaldehyde oxidation). Likewise, (iv) genes providing adaptations to oligotrophic conditions have been transmitted horizontally from indigenous pelagic microbes (e.g., rhodopsins). The second habitat transition across the freshwater-marine boundary did not result in further genome-streamlining, but is characterized by adaptations to higher salinities acquired by HGT. ‘*Ca*. M. turicensis’ was identified as transitional taxon, retaining multiple ancestral characters while also gaining adaptations to the pelagial. In this regard, the family Methylophilaceae is an exceptional model for tracing the evolutionary history of genome-streamlining as such a collection of evolutionarily related microbes from different habitats is practically unknown for other similarly abundant genome-streamlined microbes (e.g., ‘*Ca*. Pelagibacterales’, ‘*Ca*. Nanopelagicales’).

## Acknowledgements

We thank the team of the Genetic Diversity Center Zurich (GDC) for providing sequencing facilities and help with library preparation. Thomas Posch and Eugen Loher are acknowledged for help in sampling of Lake Zurich, Petr Znachor, Pavel Rychtecký and Jiří Nedoma for help in sampling of Řimov Reservoir and Lake Medard. MMS was supported by the research grant 19-23469S (Grant Agency of the Czech Republic). RG was supported by the research grant 17-04828S (Grant Agency of the Czech Republic). Sampling for the isolation of novel strains from Lake Zurich was supported by the SNF D-A-CH project 310030E-160603/1 awarded to Thomas Posch.

## Authors’ contributions

MMS conceived the project, isolated and sequenced the strains, analysed the data and wrote the manuscript. DS, MK and SMN sequenced the strains and contributed to data analyses. RG developed programs for analysis and contributed to data analyses. All authors helped to interpret the results and contributed to writing the manuscript.

