## Supplementary Figures for "Evolution in action: Habitat-transition leads to genome-streamlining in Methylophilaceae (Betaproteobacteriales)"

**Supplementary Tables S1-S5 and a set of HMMs of can be found as separate dataset (SalcherTablesS1-S5.xlsx; Salcher_methdbHMM.gz)**

**Table S1:** Characteristics of analysed genomes.

**Table S2:** Public metagenomes used for recruitment analyses and RPKG values (reads per kb of genome per Gb of metagenome) of all genomes.

**Table S3:** Membrane transporters annotated in the genomes.

**Table S4:** Selected metabolic modules present in the genomes.

**Table S5:** Complete list of proteins used for creating HMMs related to methylotrophic functions. The entire set of HMMs is available as supplementary data set (Salcher_meth-HMMs.txt)

**Figure S1:** Phylogeny of 39 strains of '*Ca*. Methylopumilus spp.'.

**Figure S2:** Average nucleotide identity (ANI) of 39 strains of '*Ca*. Methylopumilus sp.'.

**Figure S3:** Average amino acid identity (AAI) of Methylophilaceae.

**Figure S4:** Average nucleotide identity (ANI) of Methylophilaceae.

**Figure S5:** Genomic statistics of Methylophilaceae.

**Figure S6:** Amino acid usage of Methylophilaceae in relation to genome size.

**Figure S7:** Core- and pangenome analysis of Methylophilaceae.

**Figure S8:** Methylcitric acid (MCA) pathway.

**Figure S9:** Conserved methylotrophic pathways - H4F, methanol oxidation and RuMP.

**Figure S10:** Conserved methylotrophic pathways - methanol and formate oxidation.

**Figure S11:** XoxF and MxaF type methanol dehydrogenases.

**Figure S12:** Horizontal gene transfers of different rhodopsins.

**Figure S13:** Horizontal gene transfer of the NQR pathway in marine HTCC2181.

**Figure S14:** Horizontal gene transfer of an ectoine biosynthesis pathway and a sodium:proline symporter in marine OM43

**Figure S15:** Horizontal gene transfer an oligopeptide permease in marine OM43

**Figure S16:** Horizontal gene transfer of sodium transporters in marine OM43

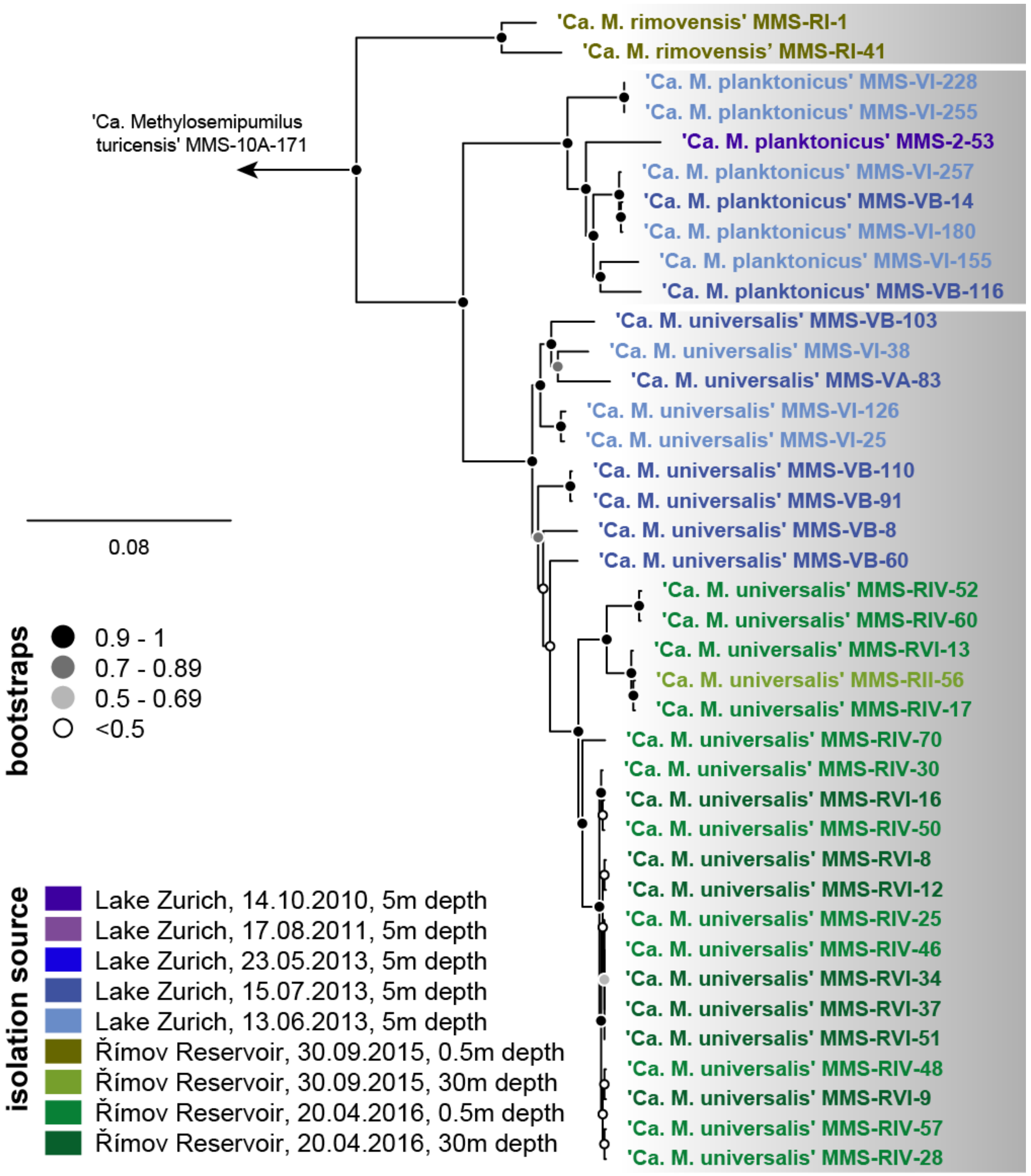


**Figure S1: Phylogeny of 39 strains of '*Ca*. Methylopumilus spp.'.**

Phylogenomic tree of 39 complete genomes of 'Ca. Methylopumilus spp' that form three distinct species ('Ca. M. rimovensis', 'Ca. M. planktonicus', 'Ca. M. universalis'). The tree is based on 983 common, concatenated protein sequences (337501 amino acid sites) with 'Ca. Methylosemipumilus turicensis' used as outgroup. Bootstraps values (100 bootstraps) are indicated by circles at individual nodes, the scale bar applies to 8% sequence divergence. The isolation source is indicated by different colours.

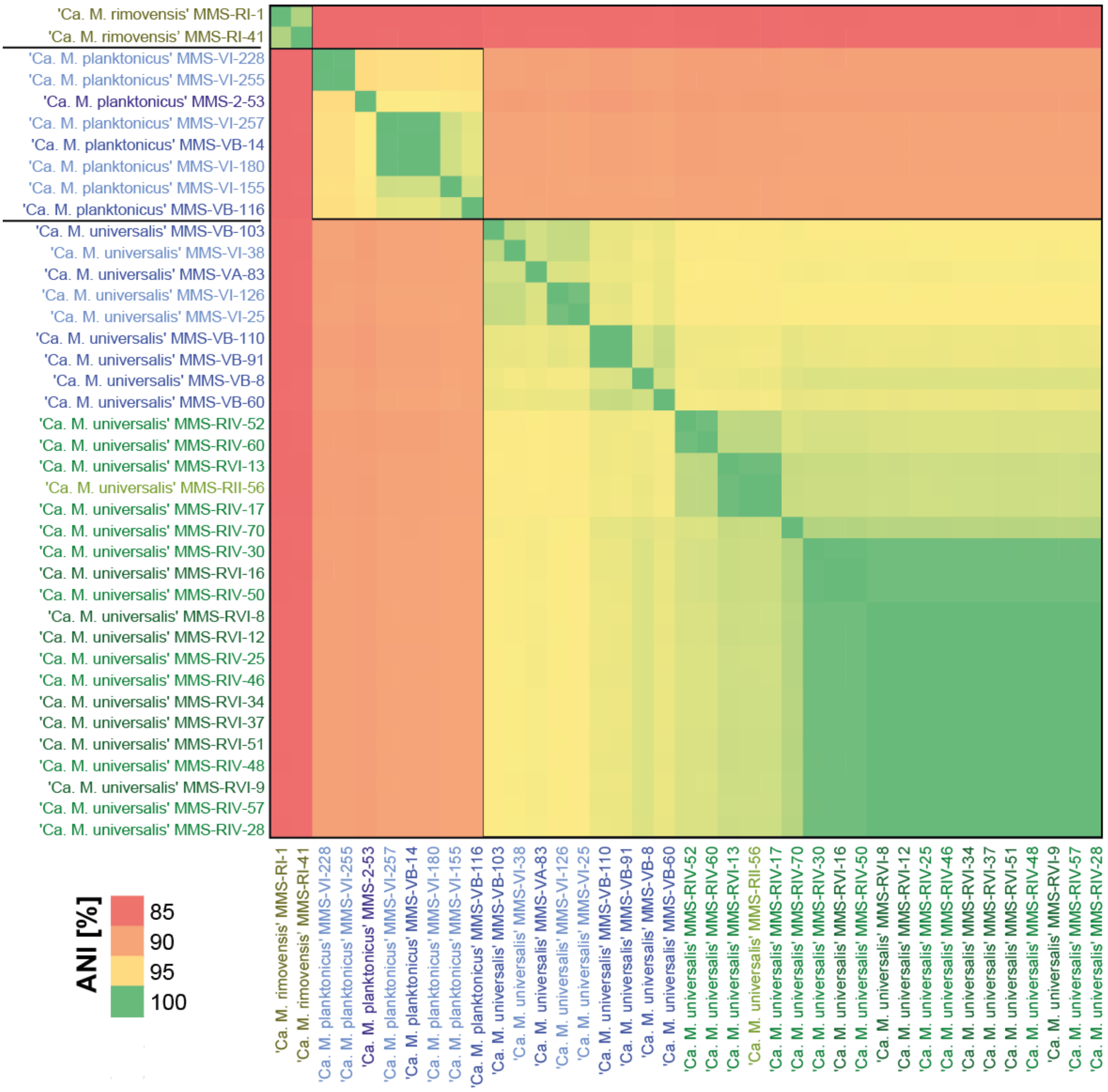


**Figure S2: Average nucleotide identity (ANI) of 39 strains of '*Ca*. Methylopumilus sp.'.**

Values >95% identity (species level delineation) are marked by black boxes. The colour coding of genomes is according to isolation source as displayed in Fig. S1.

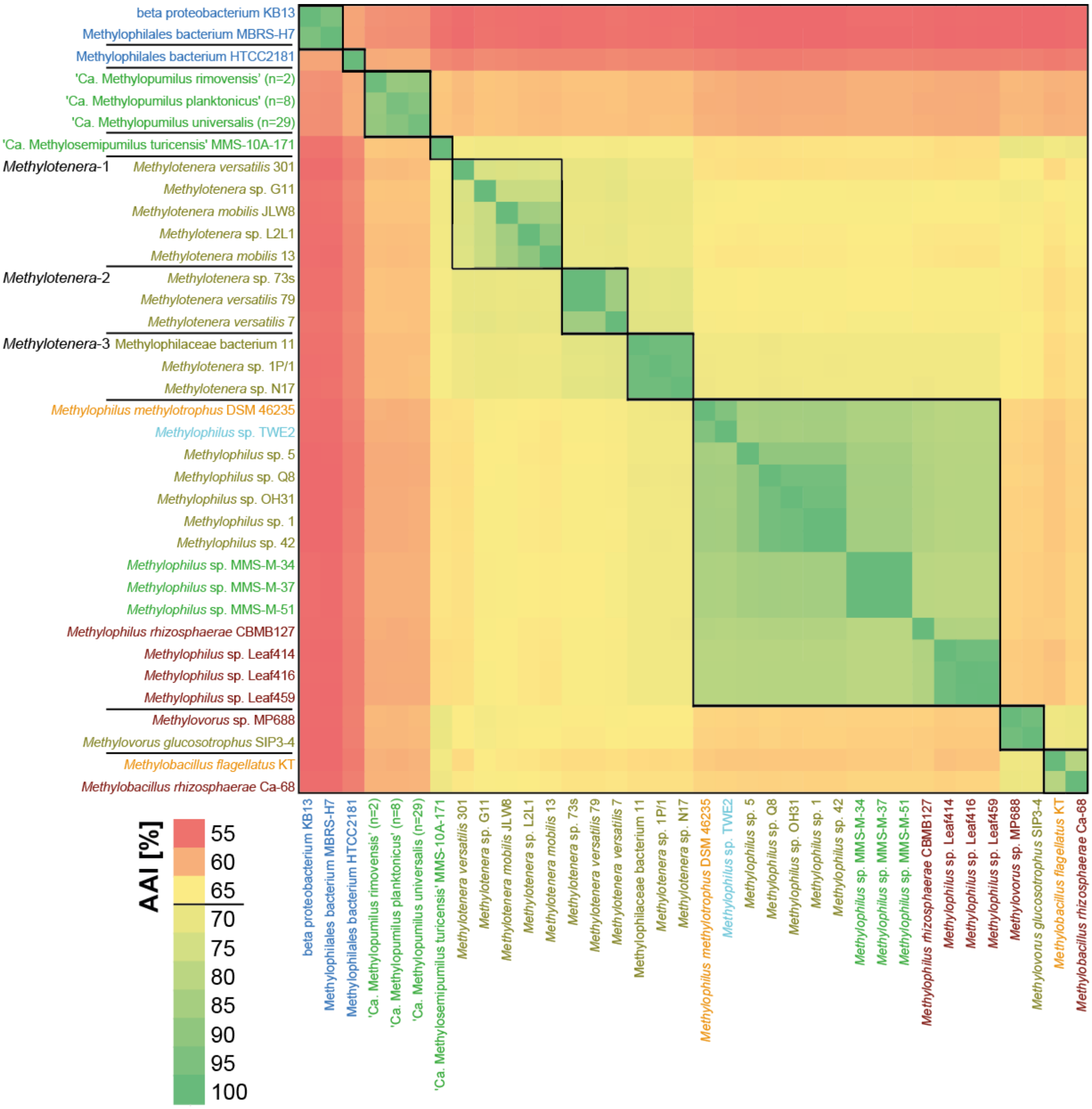


**Figure S3: Average amino acid identity (AAI) of Methylophilaceae.**

Values >70% identity (approx. genus level delineation) are marked by black boxes. The colour coding of genomes is according to Fig. 1.

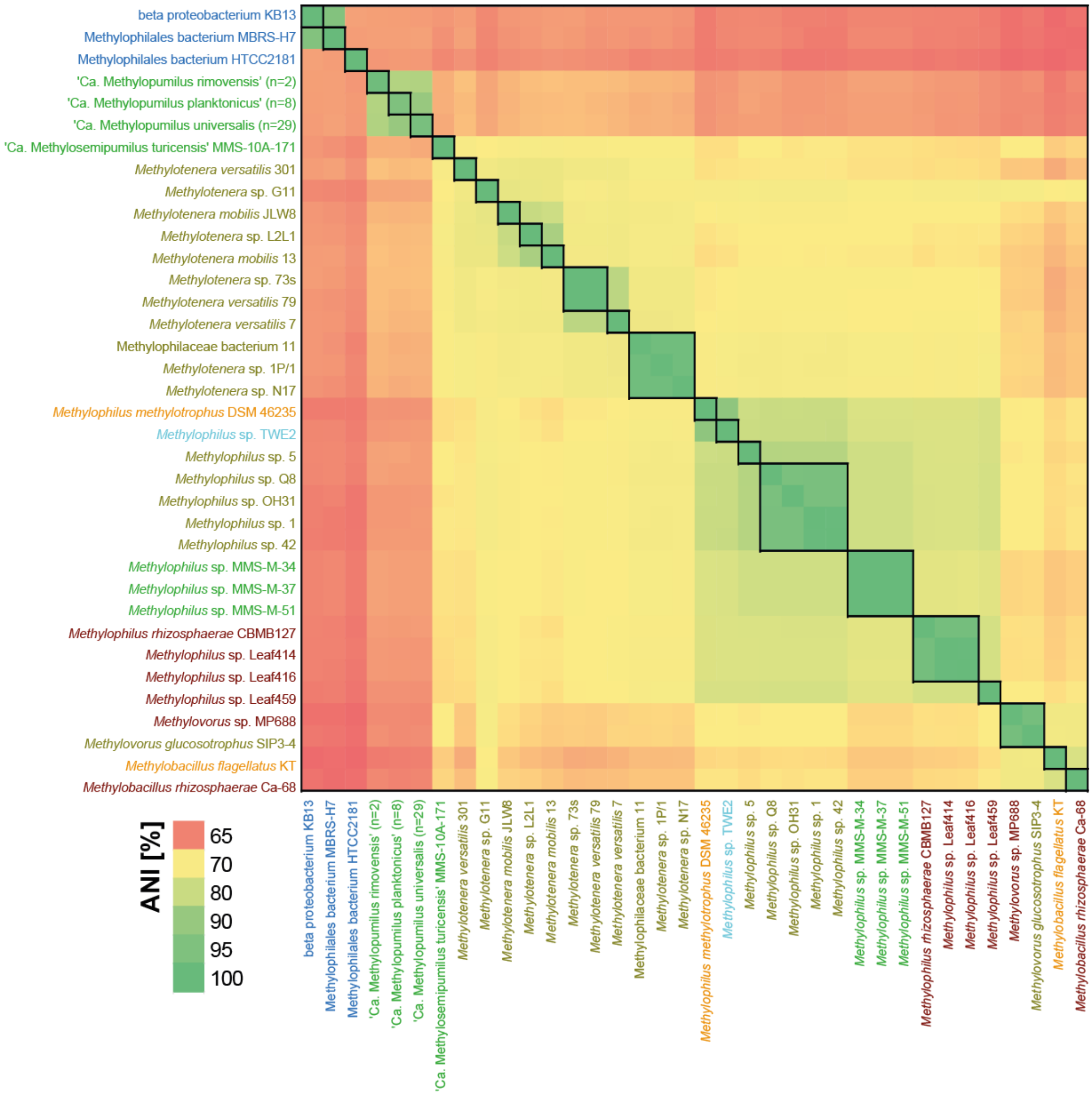


**Figure S4: Average nucleotide identity (ANI) of Methylophilaceae.**

Values >95% identity (species level delineation) are marked by black boxes. The colour coding of genomes is according to Fig. 1.

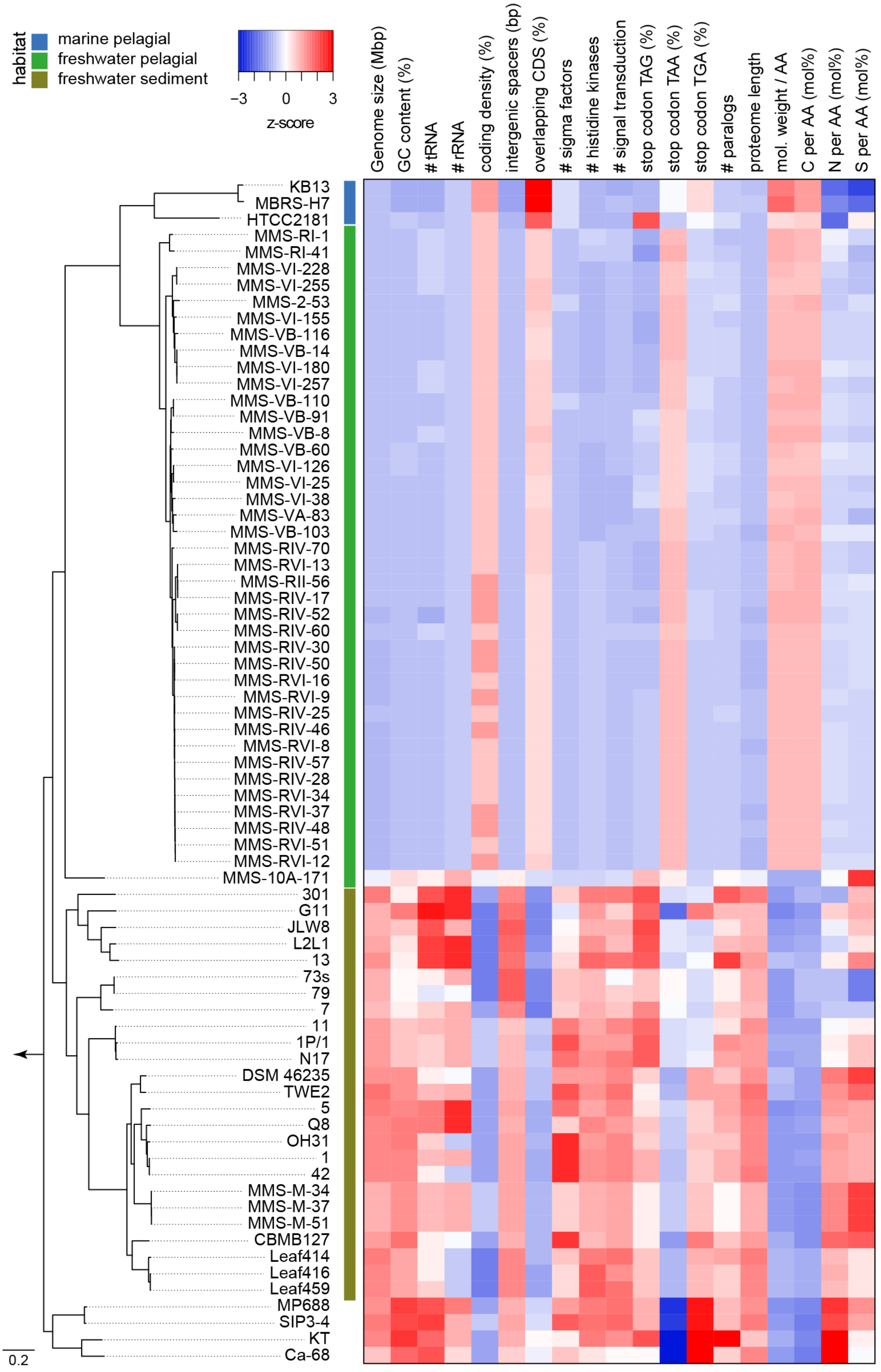


**Figure S5: Genomic statistics of Methylophilaceae.**

Phylogenomic tree (for details see Fig. 1 and Fig. S1) with selected genomic features related to genome-streamlining in Methylophilaceae.

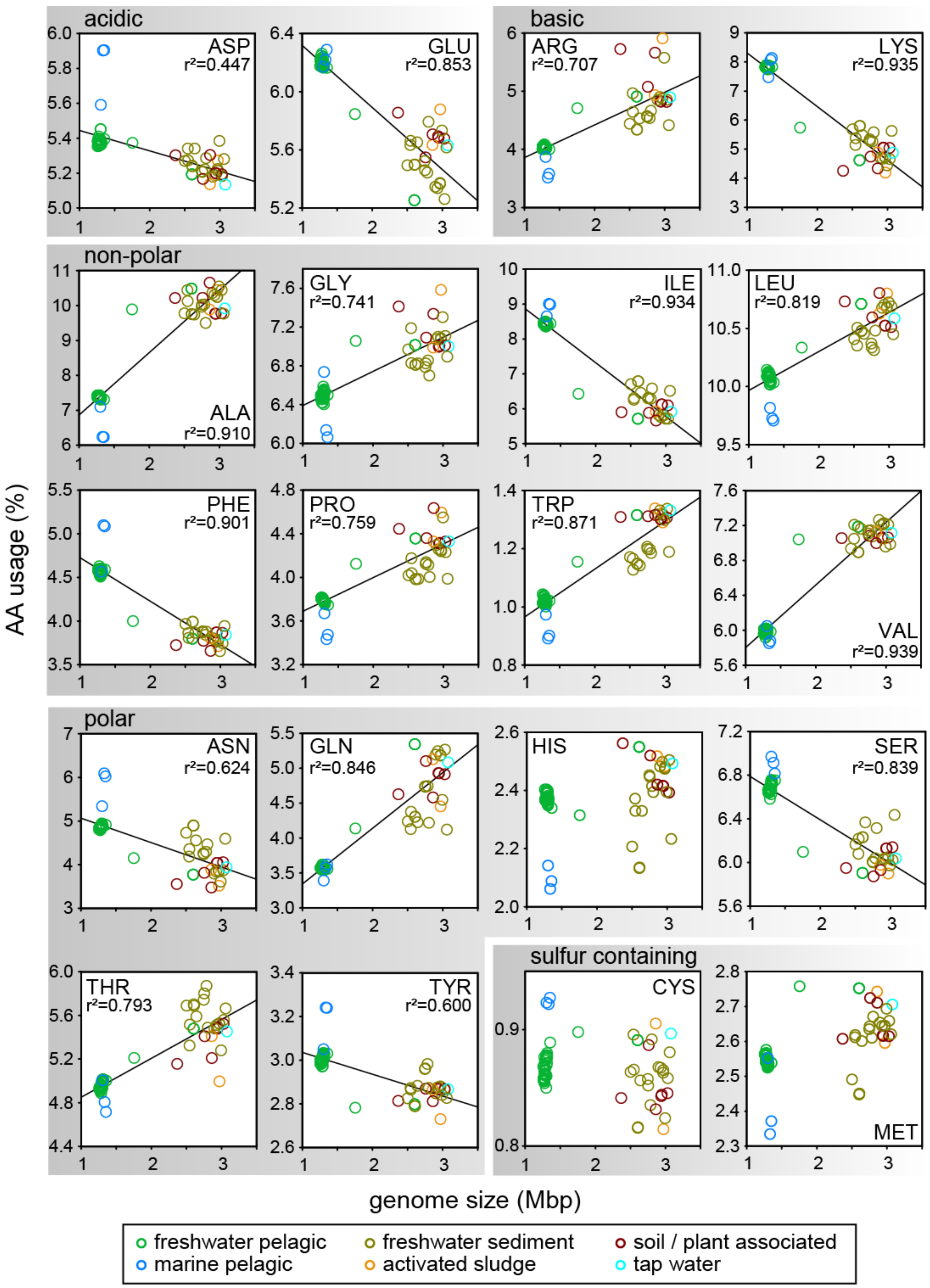


**Figure S6: Amino acid usage of Methylophilaceae in relation to genome size.**

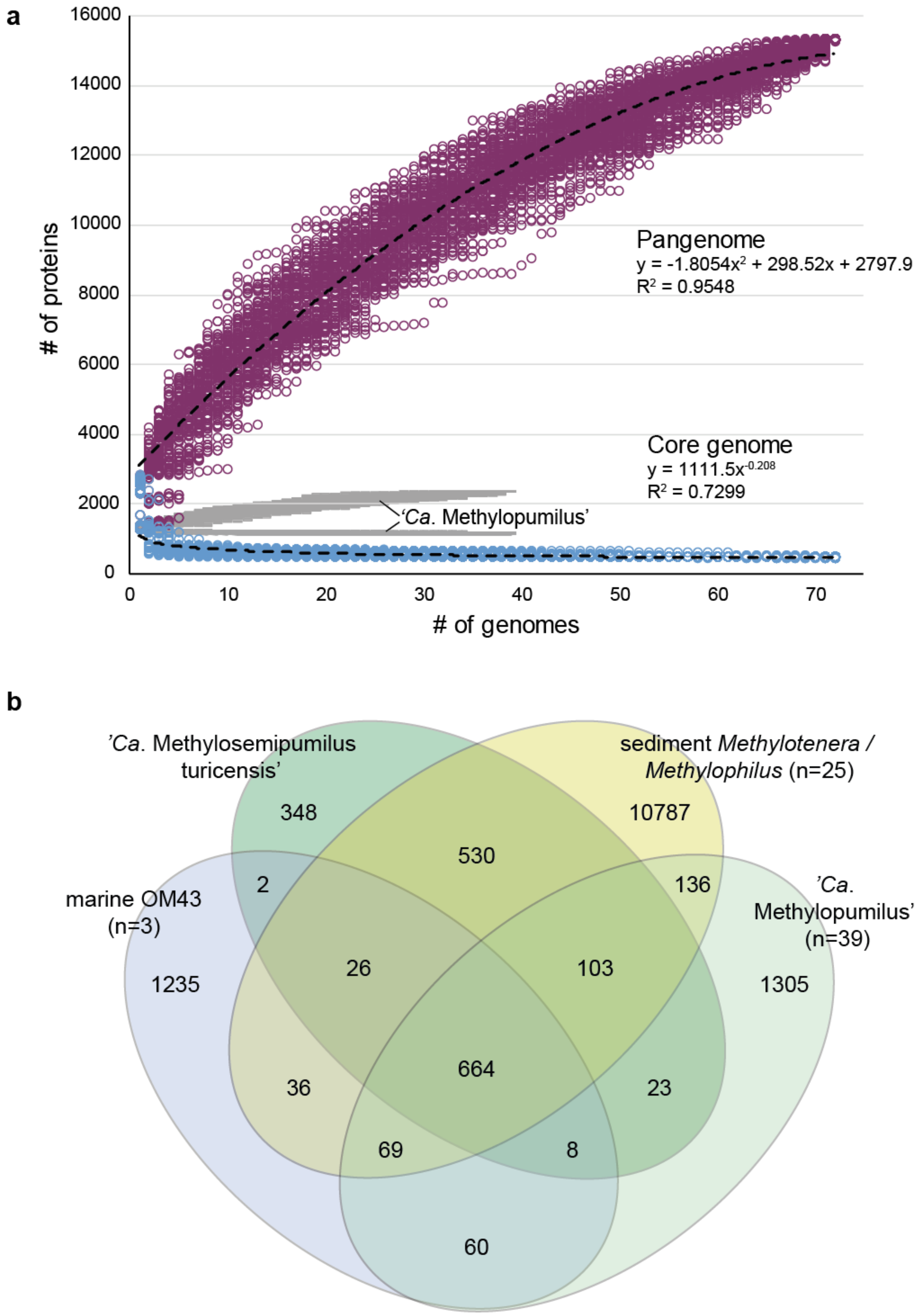


**Figure S7: Core- and pangenome analysis of Methylophilaceae.**

(a) Core- and pangenome analysis of all 72 Methylophilaceae strains in relation to 39 '*Ca*. Methylopumilus spp. strains. (b) Shared protein families of sediment *Methylophilus* and *Methylotenera*, '*Ca*. Methylosemipumilus turicensis', '*Ca*. Methylopumilus spp.' and marine OM43 at a 50% identity clustering.

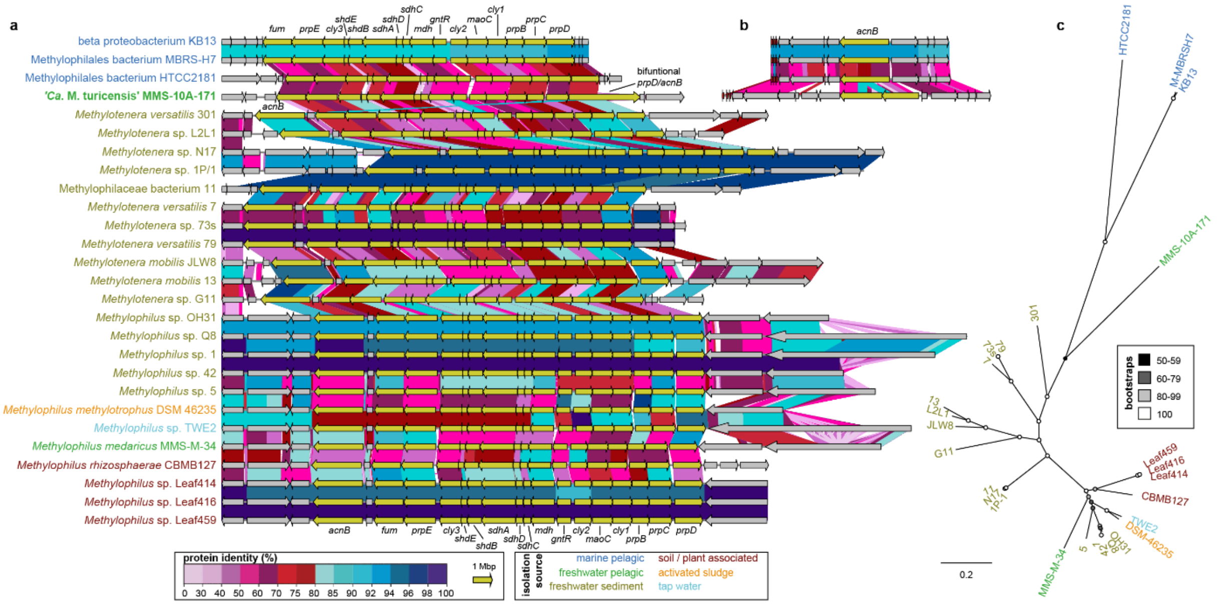


### Figure S8: Methylcitric acid (MCA) pathway

(a) Arrangement and protein similarities of genes coding for the methylcitric acid (MCA) pathway marked in dark yellow and flanking genes. (b) Similarities of *acnB* genes in marine OM43 and ‘Ca. M. turicensis’, located at different genomic regions. (c) RAxML tree (100 bootstraps) of concatenated MCA proteins.

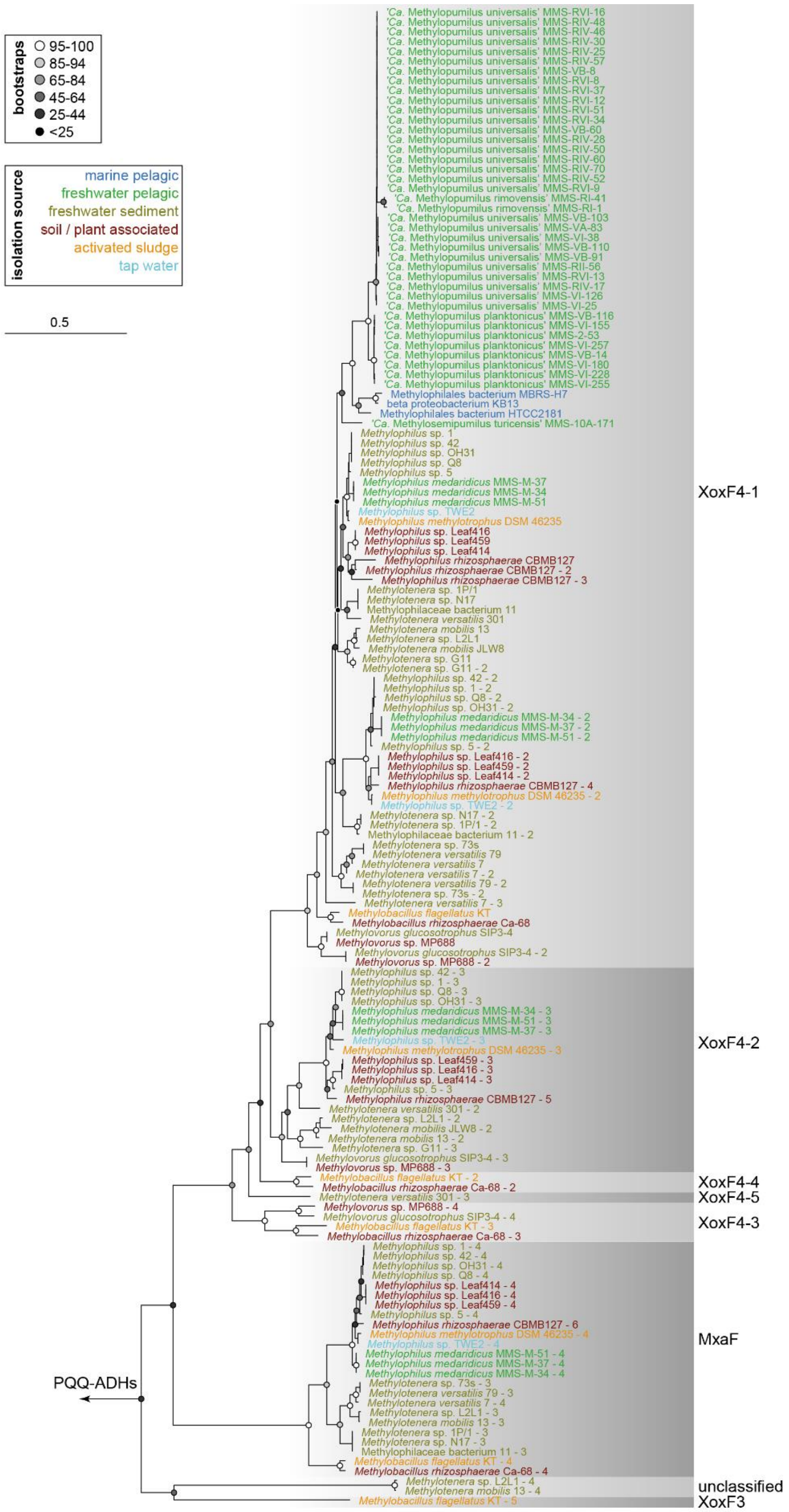


**Figure S9: XoxF and MxaF type methanol dehydrogenases**

Phylogenetic tree (100 bootstraps) of XoxF and MxaF type methanol dehydrogenases in Methylophilaceae. Isolation sources of strains are indicated by different colours. Note that some strains have multiple copies of XoxF and MxaF. Several PQQ-dependent alcohol dehydrogenases (PQQ-ADHs type 9) and other distantly related dehydrogenases were used as outgroup.

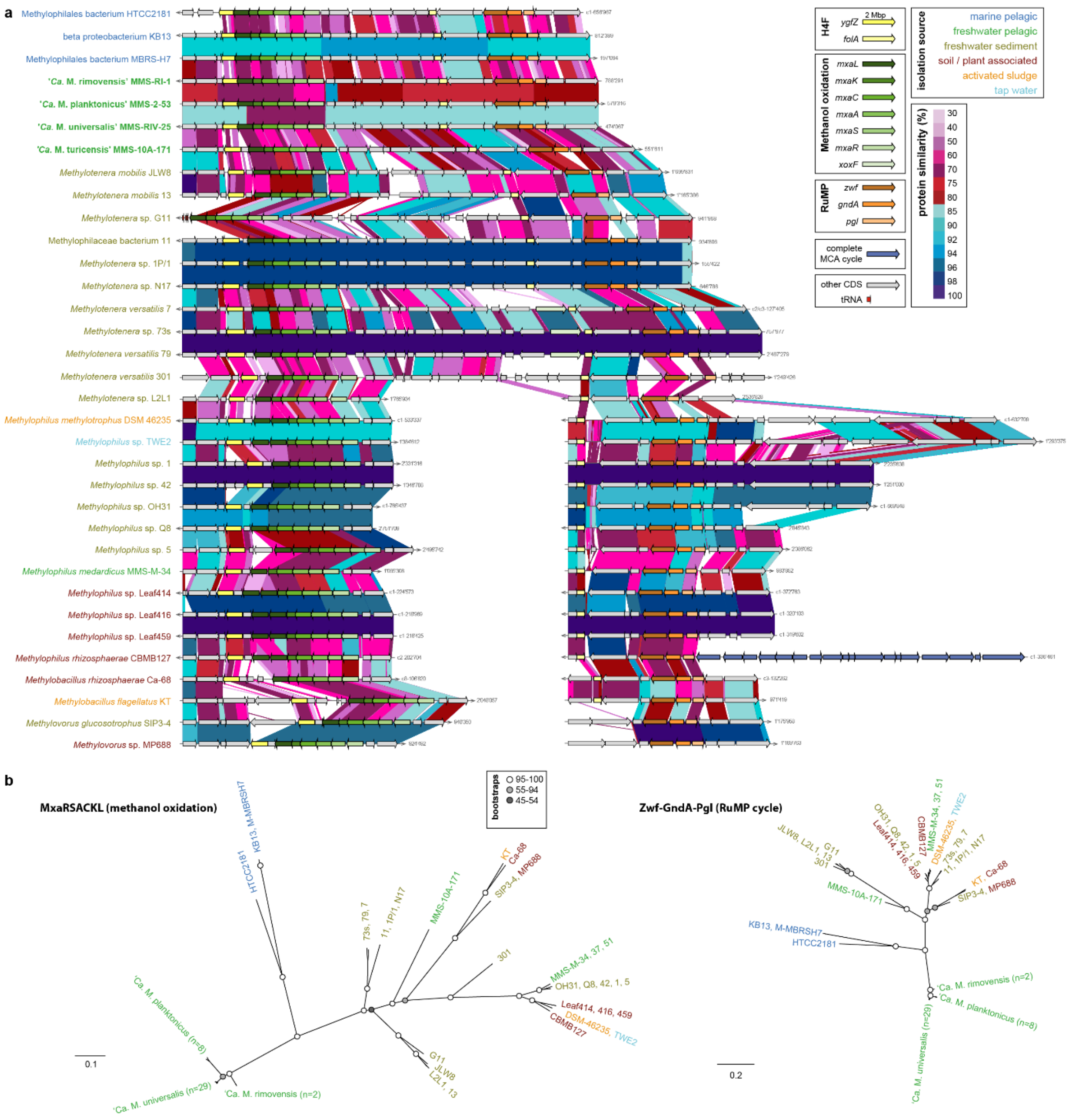

**Figure S10: Conserved methylotrophic pathways - H4F, methanol oxidation and RuMP**

(a) Synteny of different methylotrophic pathways in Methylophilaceae. Genes involved in methanol oxidation, the H4F-pathway of formaldehyde oxidation, the RuMP cycle, and a complete MCA cycle are marked by different colours, as well as protein similarities (%) of individual genes. (b) Phylogenetic trees (100 bootstraps) of concatenated MxaRSACKL (left) and Zwf-GndA-Pgl proteins (right). Isolation sources of strains are indicated by different colours.

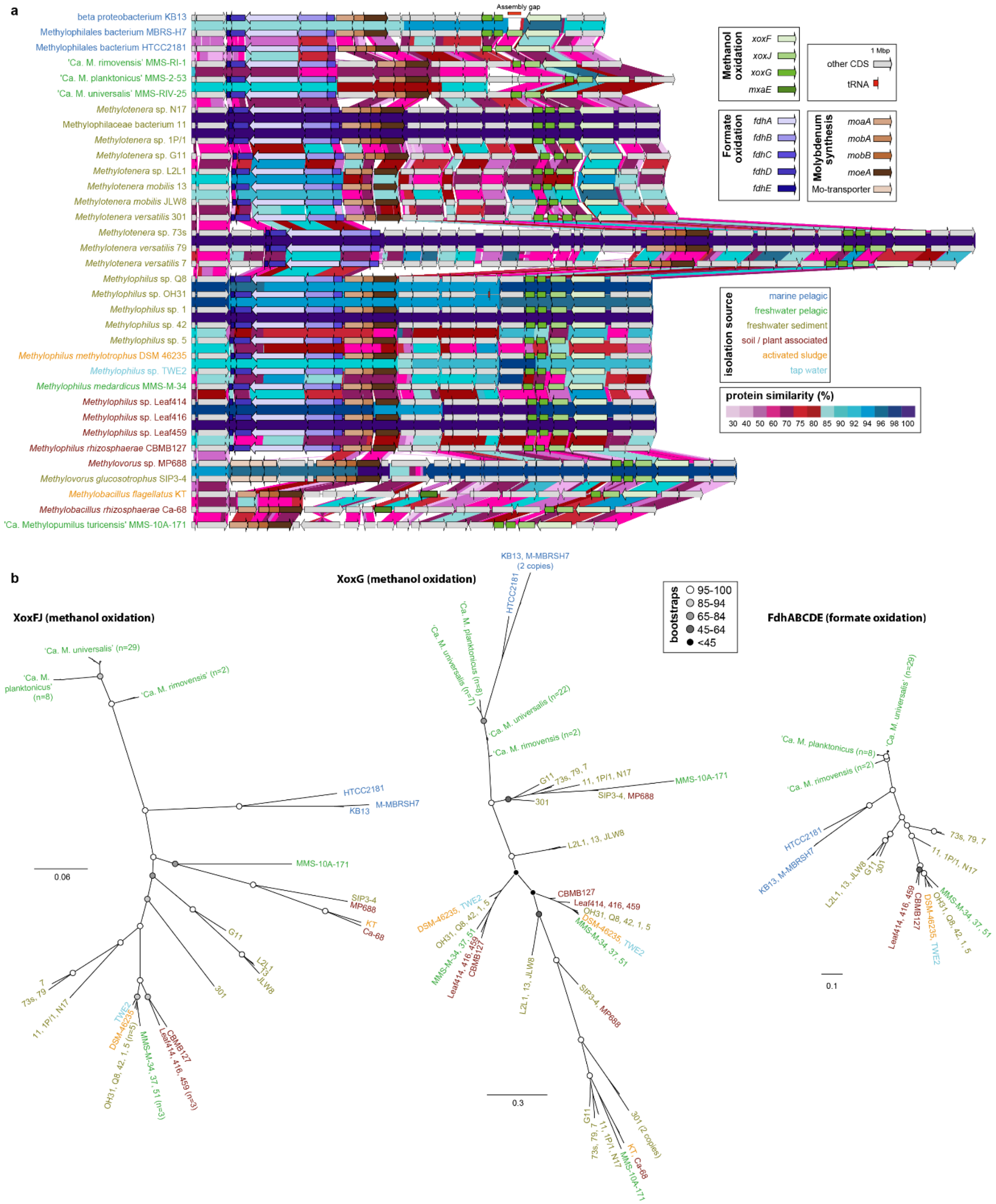


**Figure S11: Conserved methylotrophic pathways - methanol and formate oxidation**

(a) Synteny of different methylotrophic pathways in Methylophilaceae. Genes involved in methanol oxidation, formate oxidation and synthesis of the cofactor molybdenum are marked by different colours, as well as protein similarities (%) of individual genes. (b) Phylogenetic trees (100 bootstraps) of concatenated XoxFJ (left), XoxG (middle) and FdhABCDE (right). Isolation sources of strains are indicated by different colours. Note that some strains have multiple copies of XoxG.

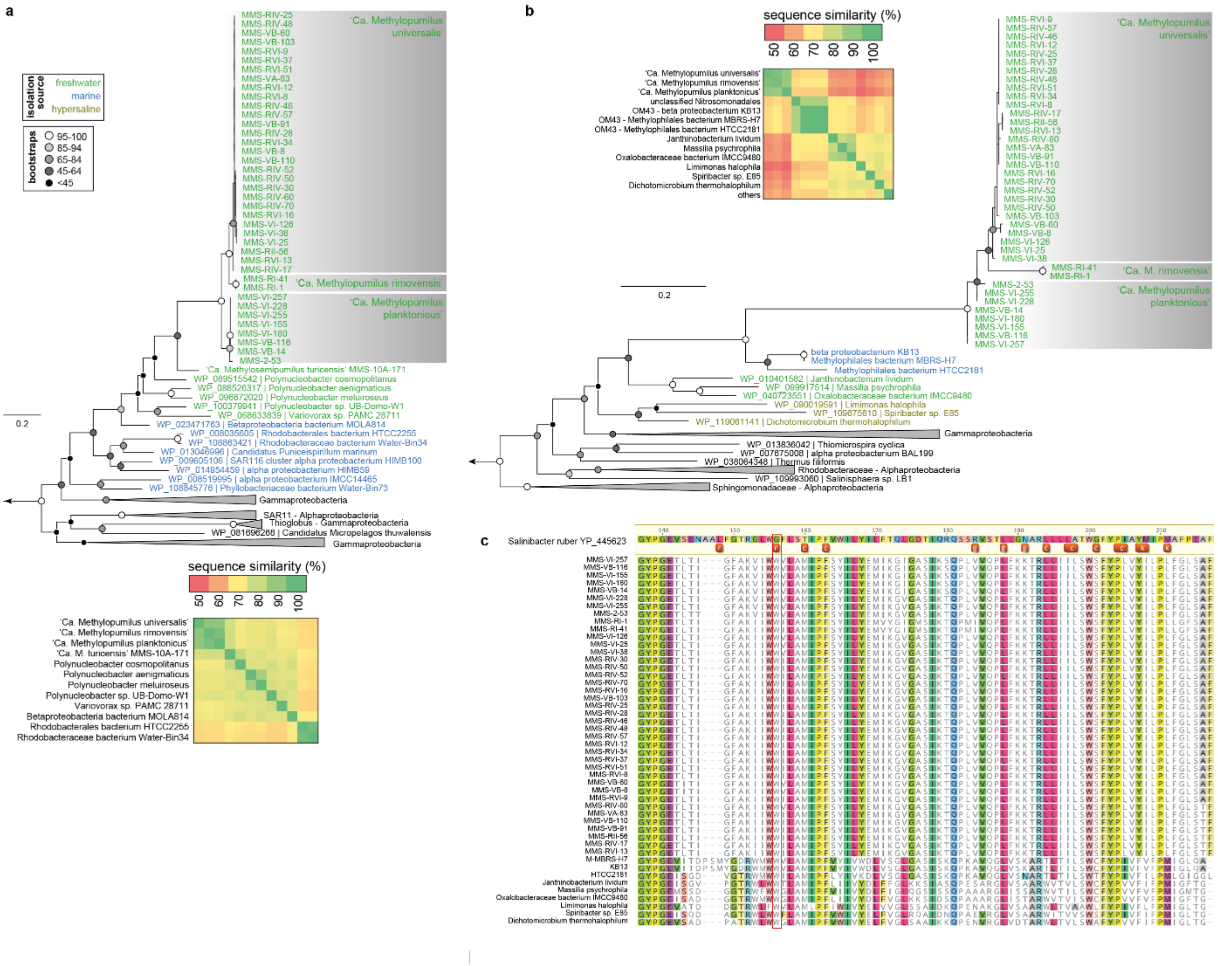


**Figure S12: Horizontal gene transfers of different rhodopsins**

RAxML trees (100 bootstraps) of (a) proteorohodopsins of '*Ca*. Methylopumilus spp.' and '*Ca*. Methylosemipumilus turicensis' and (b) xantho-like rhodopsins of '*Ca*. Methylopumilus spp.' and marine OM43. Protein sequence identities for the closest relatives are given below (a) or at the left side (b). Isolation sources of the closest relatives are indicated by different colours, actinorhodopsins were used as outgroup for both trees. (c) Amino acid alignment of xantho-like rhodopsins with residues involved in caroteinoid binding in *Salinibacter ruber* DSM13855 (Luecke et al. 2008, Imasheva et al. 2009). Letters c, g, k, and r stand for the chain, glucoside, keto group, and ring, respectively, and indicate part of the carotenoid in the vicinity of a residue. Note ring binding site at position 156 highlighted in red, where glycine is replaced with tryptophan in all strains indicating that the antenna might not bind.

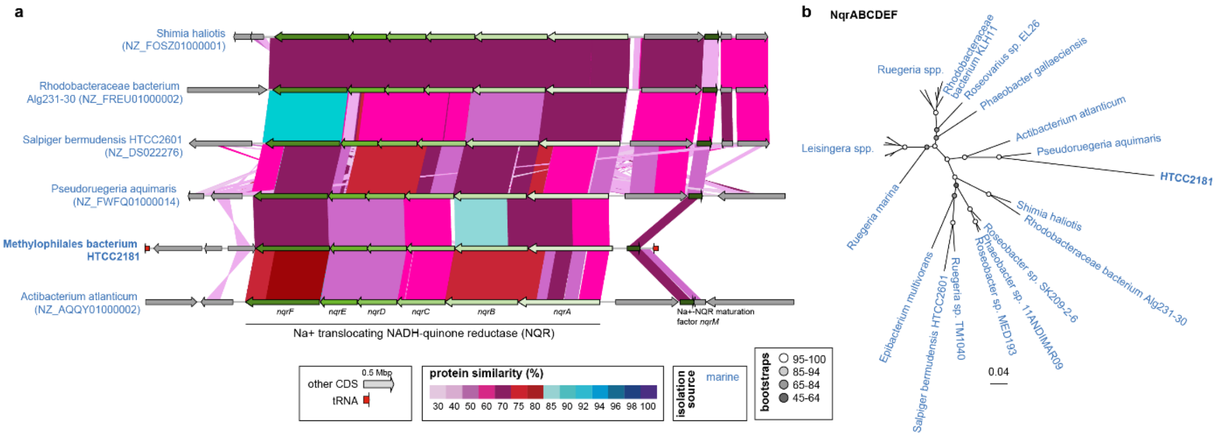


**Figure S13: Horizontal gene transfer of the NQR pathway in marine HTCC2181**

(a) Arrangement and protein similarities of genes coding for the Na+ translocating NADH-quinone reductase (NQR) marked in green colours and flanking genes. (b) RAxML tree (100 bootstraps) of concatenated NqrABCDEF proteins of the NQR pathway. Isolation sources of the strains are indicated by different colours.

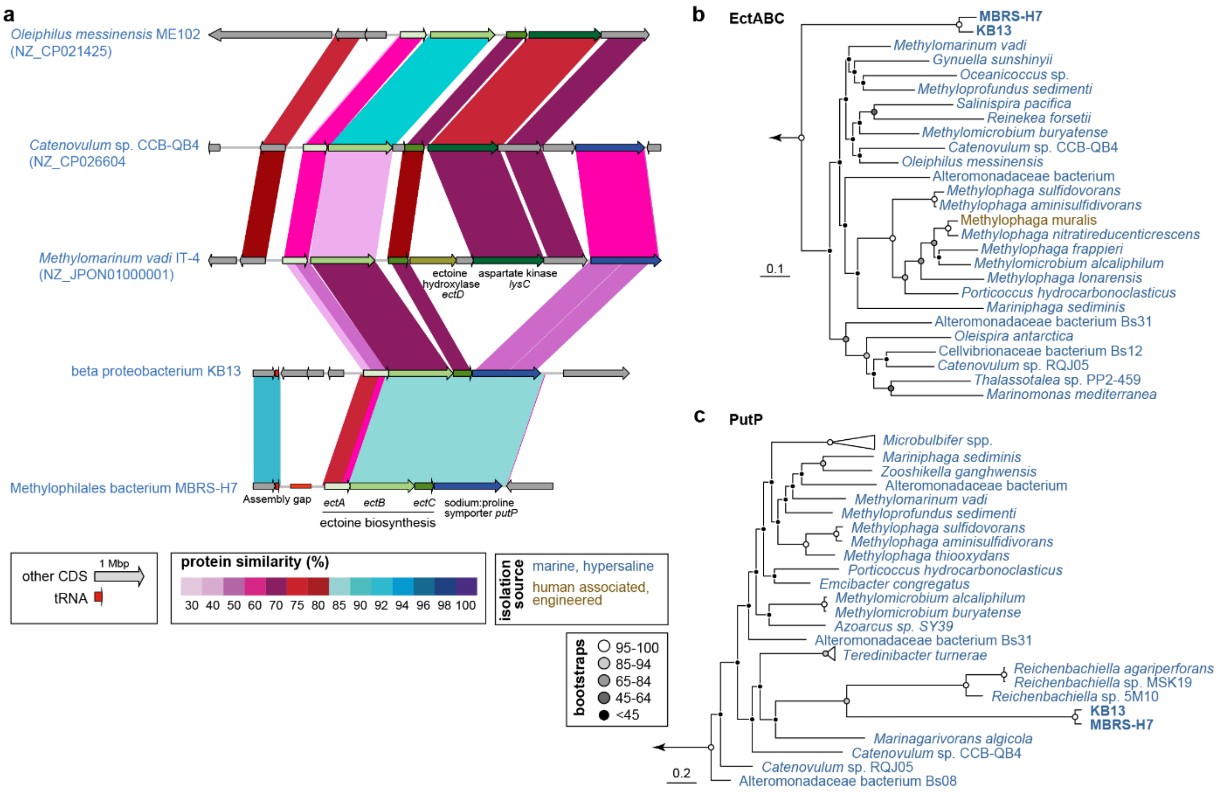


**Figure S14: Horizontal gene transfer of an ectoine biosynthesis pathway and a sodium:proline symporter in marine OM43**

(a) Arrangement and protein similarities of genes coding for the ectoine biosynthesis pathway (*ectABC*) marked in green colours and flanking genes, including a sodium:proline symporter (marked in blue). Aspartate kinase *lysC* as well as aspartate-semialdehyde dehydrogenase *asd* are located elsewhere in the genomes of KB13 and MBRS-H7 and more similar to *lysC* and *asd* of other Methylophilaceae. RAxML trees (100 bootstraps) of (b) concatenated EctABC proteins of the ectoine biosynthesis pathway and (c) the sodium:proline symporter PutP. Isolation sources of the strains are indicated by different colours.

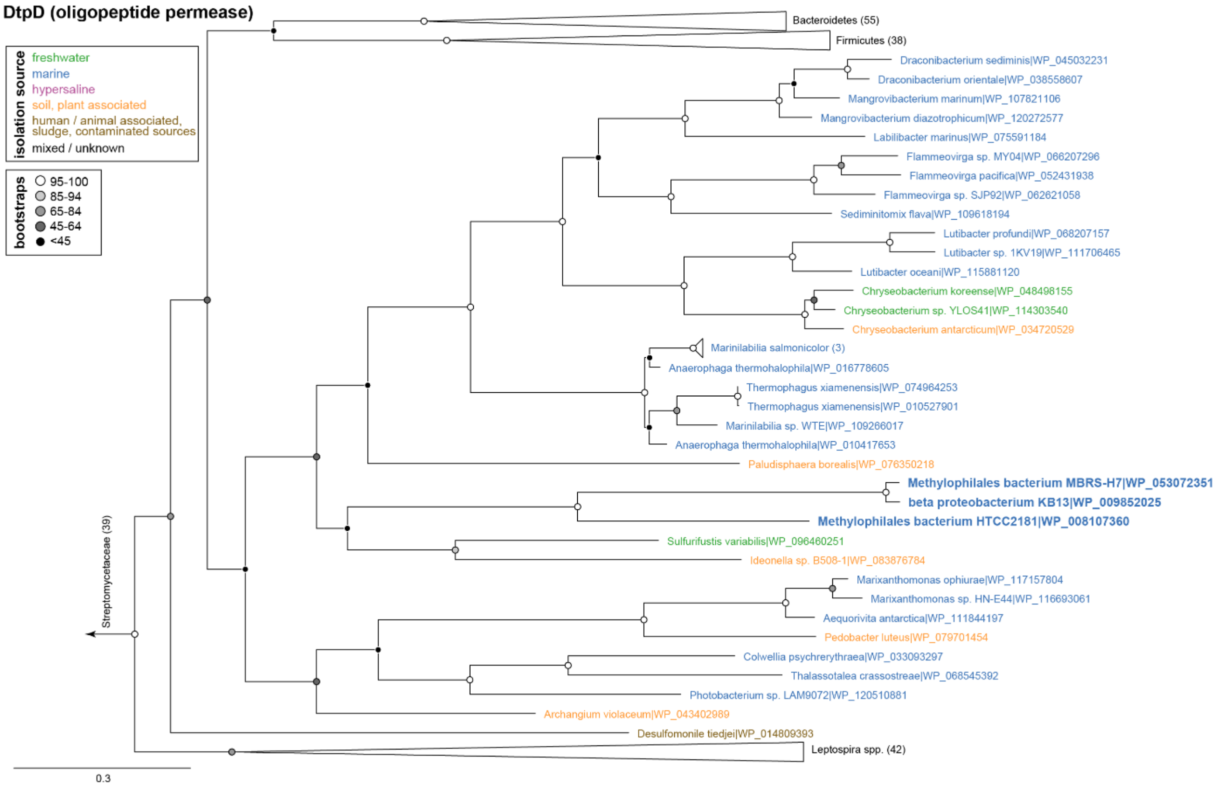


**Figure S15: Horizontal gene transfer an oligopeptide permease in marine OM43**

RAxML trees (100 bootstraps) of the dipeptide/tripeptide permease DtpD unique for the marine OM43 cluster (in bold). Isolation sources of the closest relatives are indicated in different colours.

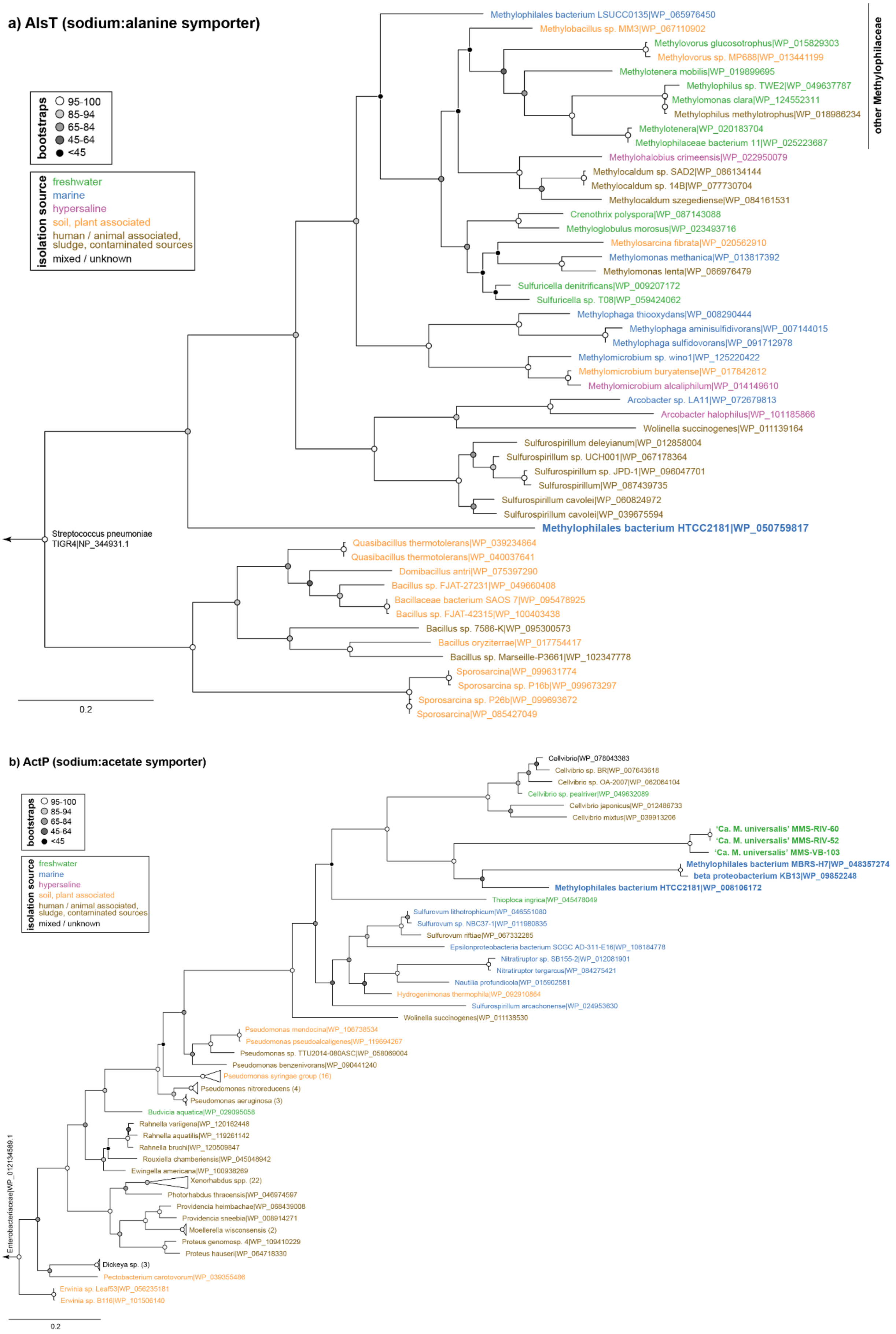

a) AlsT (sodium:alanine symporter)
bootstraps
95-100
85-94
65-84
45-64
<45
isolation source
freshwater
marine
hypersaline
soil, plant associated
human / animal associated, sludge, contaminated sources
mixed / unknown
other Methylophilaceae
Methylophilales bacterium LSUCC0135|WP_065976450
Methylobacillus sp. MM3|WP_067110902
Methylovorus glucosotrophus|WP_015829303
Methylovorus sp. MP688|WP_013441199
Methylotenera mobilis|WP_019899695
Methylophilus sp. TWE2|WP_049637787
Methylomonas clara|WP_124552311
Methylophilus methylotrophus|WP_018986234
Methylotenera|WP_020183704
Methylophilaceae bacterium 11|WP_025223687
Methylohalobius crimeensis|WP_022950079
Methylocaldum sp. SAD2|WP_086134144
Methylocaldum sp. 14B|WP_077730704
Methylocaldum szegediense|WP_084161531
Crenothrix polyspora|WP_087143088
Methyloglobulus morosus|WP_023493716
Methylosarcina fibrata|WP_020562910
Methylomonas methanica|WP_013817392
Methylomonas lenta|WP_066976479
Sulfuricella denitrificans|WP_009207172
Sulfuricella sp. T08|WP_059424062
Methylophaga thiooxydans|WP_008290444
Methylophaga aminisulfidivorans|WP_007144015
Methylophaga sulfidovorans|WP_091712978
Methylomicrobium sp. wino1|WP_125220422
Methylomicrobium buryatense|WP_017842612
Methylomicrobium alcaliphilum|WP_014149610
Arcobacter sp. LA11|WP_072679813
Arcobacter halophilus|WP_101185866
Wolinella succinogenes|WP_011139164
Sulfurospirillum deleyianum|WP_012858004
Sulfurospirillum sp. UCH001|WP_067178364
Sulfurospirillum sp. JPD-1|WP_096047701
Sulfurospirillum|WP_087439735
Sulfurospirillum cavolei|WP_060824972
Sulfurospirillum cavolei|WP_039675594
Methylophilales bacterium HTCC2181|WP_050759817
Streptococcus pneumoniae TIGR4|NP_344931.1
Quasibacillus thermotolerans|WP_039234864
Quasibacillus thermotolerans|WP_040037641
Domibacillus antri|WP_075397290
Bacillus sp. FJAT-27231|WP_049660408
Bacillaceae bacterium SAOS 7|WP_095478925
Bacillus sp. FJAT-42315|WP_100403438
Bacillus sp. 7586-K|WP_095300573
Bacillus oryziterrae|WP_017754417
Bacillus sp. Marseille-P3661|WP_102347778
Sporosarcina|WP_099631774
Sporosarcina sp. P16b|WP_099673297
Sporosarcina sp. P26b|WP_099693672
Sporosarcina|WP_085427049
0.2
b) ActP (sodium:acetate symporter)
bootstraps
95-100
85-94
65-84
45-64
<45
isolation source
freshwater
marine
hypersaline
soil, plant associated
human / animal associated, sludge, contaminated sources
mixed / unknown
Cellvibrio|WP_078043383
Cellvibrio sp. BR|WP_007643618
Cellvibrio sp. OA-2007|WP_062064104
Cellvibrio sp. pealriver|WP_049632089
Cellvibrio japonicus|WP_012486733
Cellvibrio mixtus|WP_039913206
'Ca. M. universalis' MMS-RIV-60
'Ca. M. universalis' MMS-RIV-52
'Ca. M. universalis' MMS-VB-103
Methylophilales bacterium MBRS-H7|WP_048357274
beta proteobacterium KB13|WP_09852248
Methylophilales bacterium HTCC2181|WP_008106172
Thioploca ingrica|WP_045478049
Sulfurovum lithotrophicum|WP_046551080
Sulfurovum sp. NBC37-1|WP_011980835
Sulfurovum riftiae|WP_067332285
Epsilonproteobacteria bacterium SCGC AD-311-E16|WP_106184778
Nitratiruptor sp. SB155-2|WP_012081901
Nitratiruptor tergarcus|WP_084275421
Nautilia profundicola|WP_015902581
Hydrogenimonas thermophila|WP_092910864
Sulfurospirillum arcachonense|WP_024953630
Wolinella succinogenes|WP_011138530
Pseudomonas mendocina|WP_106738534
Pseudomonas pseudoalcaligenes|WP_119694267
Pseudomonas sp. TTU2014-080ASC|WP_058069004
Pseudomonas benzenivorans|WP_090441240
Pseudomonas syringae group (16)
Pseudomonas nitroreducens (4)
Pseudomonas aeruginosa (3)
Budvicia aquatica|WP_029095058
Rahnella variigena|WP_120162448
Rahnella aquatilis|WP_119261142
Rahnella bruchi|WP_120509847
Rouxiella chamberiensis|WP_045048942
Ewingella americana|WP_100938269
Xenorhabdus spp. (22)
Photorhabdus thracensis|WP_046974597
Providencia heimbachae|WP_068439008
Providencia sneebia|WP_008914271
Moellerella wisconsensis (2)
Proteus genomosp. 4|WP_109410229
Proteus hauseri|WP_064718330
Dickeya sp. (3)
Pectobacterium carotovorum|WP_039355486
Erwinia sp. Leaf53|WP_056235181
Erwinia sp. B116|WP_101506140
Enterobacteriaceae|WP_012134589.1
0.2

#### c) GltT (sodium:dicarboxylate symporter)

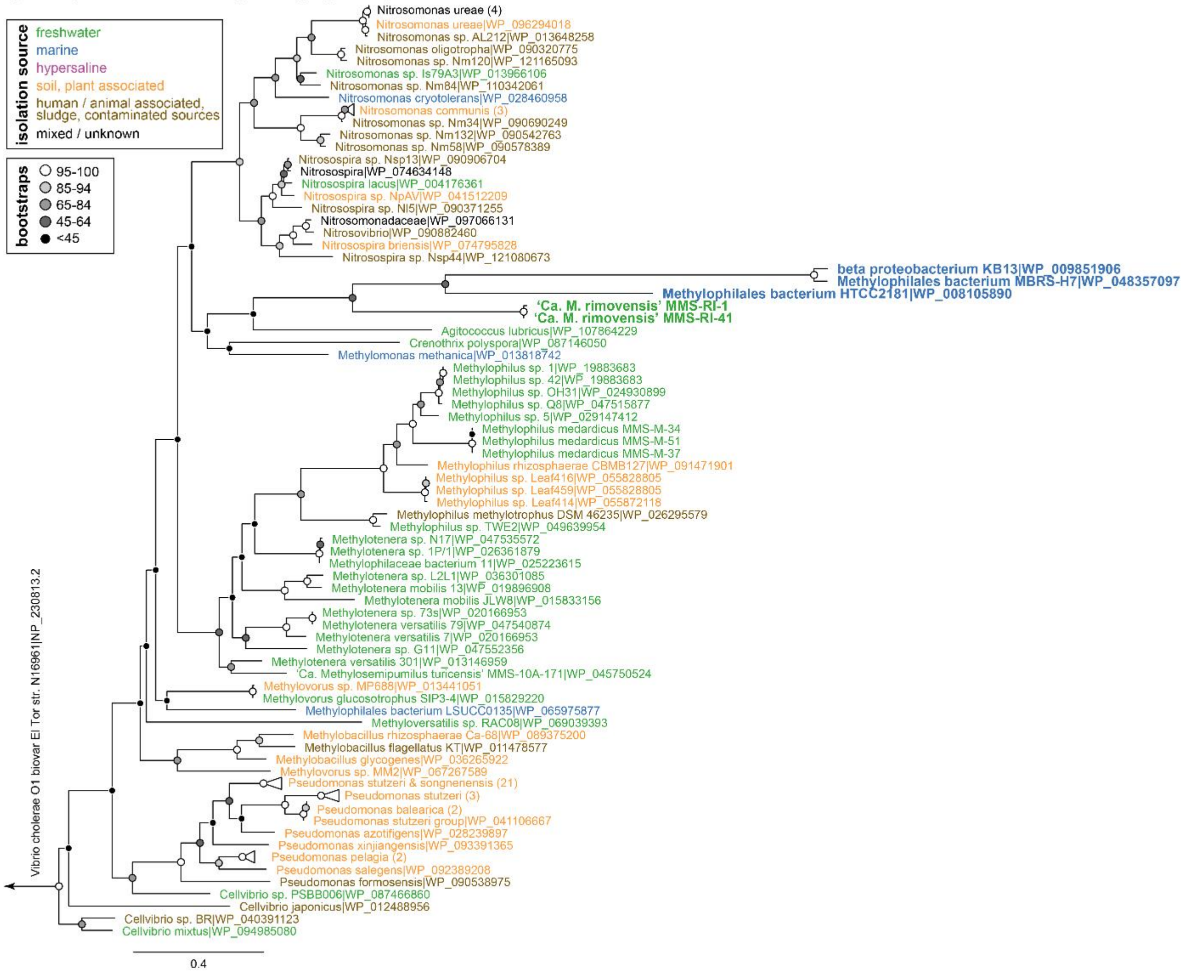


#### d) NhaP2 (sodium:proton antiporter)

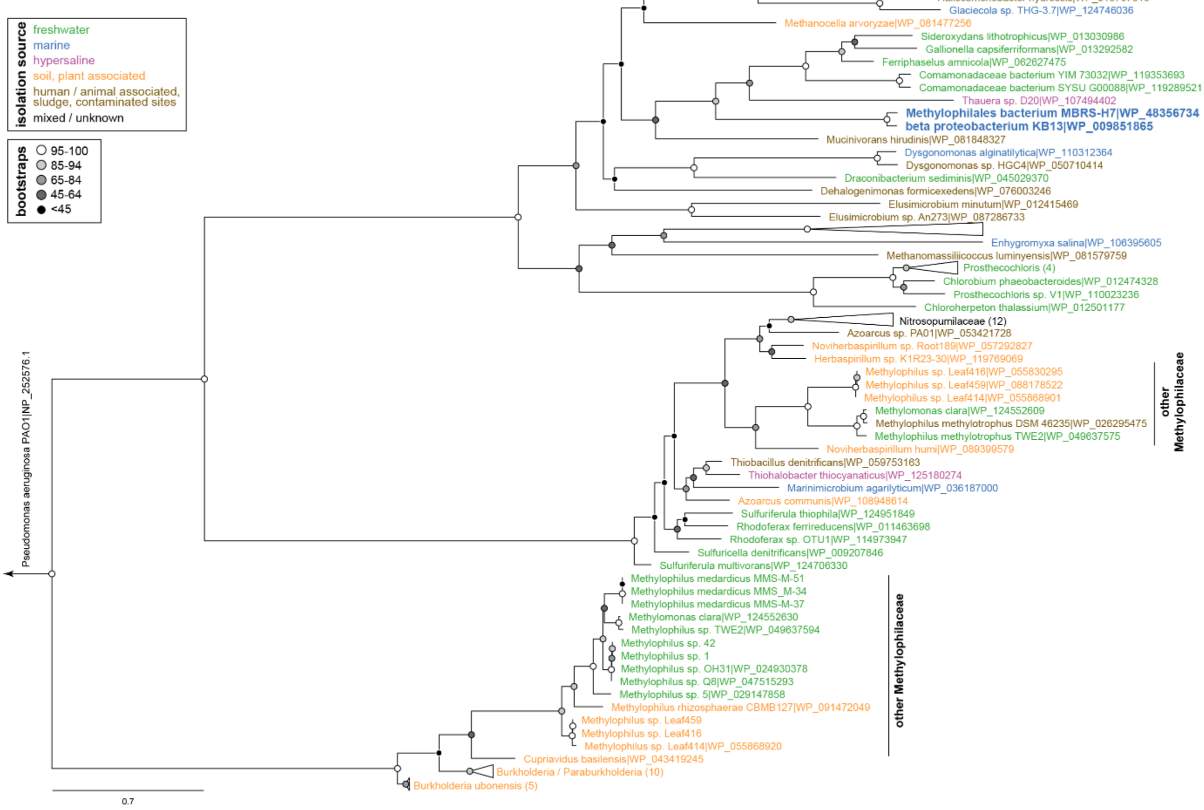

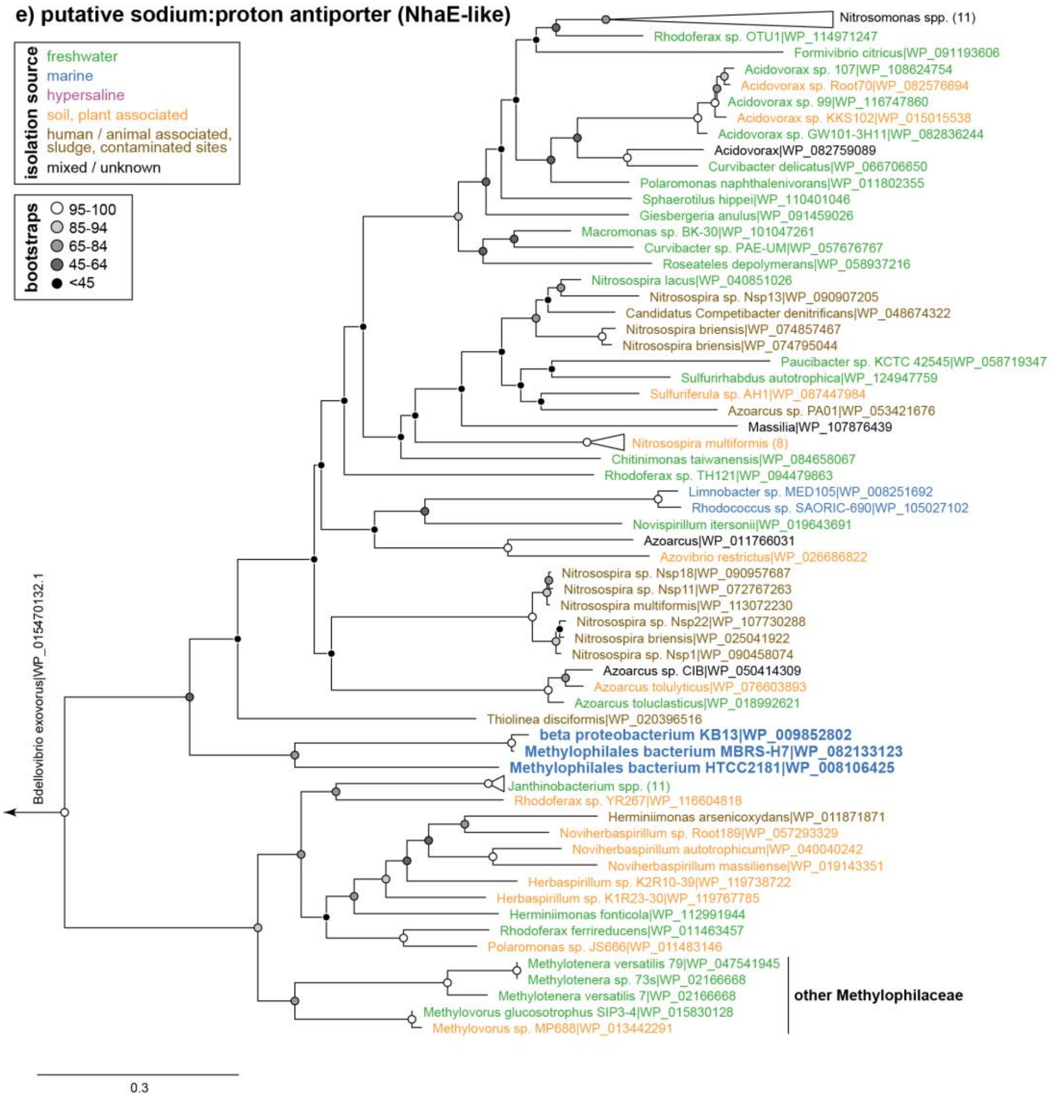


**Figure S16: Horizontal gene transfer of sodium transporters in marine OM43**

RAxML trees (100 bootstraps) of (a) the sodium:alanine symporter AlsT, the sodium:acetate symporter ActP, (c) the sodium:dicarboxylate symporter GltT, (d) the sodium:proton antiporter NhaP2, and (e) another putative sodium:proton antiporter (NhaE-like). Marine OM43 and ‘*Ca*. Methylopumilus‘ strains are highlighted in bold, isolation sources of the closest relatives are indicated in different colours.
